# Modeling the growth of organisms validates a general relation between metabolic costs and natural selection

**DOI:** 10.1101/358440

**Authors:** Efe Ilker, Michael Hinczewski

## Abstract

Metabolism and evolution are closely connected: if a mutation incurs extra energetic costs for an organism, there is a baseline selective disadvantage that may or may not be compensated for by other adaptive effects. A long-standing, but to date unproven, hypothesis is that this disadvantage is equal to the fractional cost relative to the total resting metabolic expenditure. This hypothesis has found a recent resurgence as a powerful tool for quantitatively understanding the strength of selection among different classes of organisms. Our work explores the validity of the hypothesis from first principles through a generalized metabolic growth model, versions of which have been successful in describing organismal growth from single cells to higher animals. We build a mathematical framework to calculate how perturbations in maintenance and synthesis costs translate into contributions to the selection coefficient, a measure of relative fitness. This allows us to show that the hypothesis is an approximation to the actual baseline selection coefficient. Moreover we can directly derive the correct prefactor in its functional form, as well as analytical bounds on the accuracy of the hypothesis for any given realization of the model. We illustrate our general framework using a special case of the growth model, which we show provides a quantitative description of overall metabolic synthesis and maintenance expenditures in data collected from a wide array of unicellular organisms (both prokaryotes and eukaryotes). In all these cases we demonstrate that the hypothesis is an excellent approximation, allowing estimates of baseline selection coefficients to within 15% of their actual values. Even in a broader biological parameter range, covering growth data from multicellular organisms, the hypothesis continues to work well, always within an order of magnitude of the correct result. Our work thus justifies its use as a versatile tool, setting the stage for its wider deployment.

## Introduction

Discovering optimality principles in biological function has been a major goal of biophysics [1–6], but the competition between genetic drift and natural selection means that evolution is not purely an optimization process [7–9]. A necessary complement to elucidating optimality is clarifying under what circumstances selection is actually strong enough relative to drift in order to drive systems toward local optima in the fitness landscape. In this work we focus on one key component of this problem: quantifying the selective pressure on the extra metabolic costs associated with a genetic variant. We validate a long hypothesized relation [10–12] between this pressure and the fractional change in the total resting metabolic expenditure of the organism.

The effectiveness of selection versus drift hinges on two non-dimensional parameters [13]: i) the *selection coefficient s*, a measure of the fitness of the mutant versus the wild-type. Mutants will have on average 1+*s* offspring relative to the wild-type per wild-type generation time; ii) the *effective population N*_*e*_ of the organism, the size of an idealized, randomly mating population that exhibits the same decrease in genetic diversity per generation due to drift as the actual population (with size *N*). For a deleterious mutant (*s <* 0) where 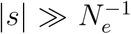, natural selection is dominant, with the probability of the mutant fixing in the population exponentially suppressed. In contrast if 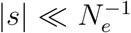, drift is dominant, with the fixation probability being approximately the same as for a neutral mutation [7]. Thus the magnitude of 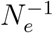 determines the “drift barrier” [14], the critical minimum scale of the selection coefficient for natural selection to play a non-negligible role.

The long-term effective population size *N*_*e*_ of an organism is typically smaller than the instantaneous actual *N*, and can be estimated empirically across a broad spectrum of life: it varies from as high as 10^9^ *-* 10^10^ in many bacteria, to 10^6^ *-* 10^8^ in unicellular eukaryotes, down to *∼* 10^6^ in invertebrates and *∼* 10^4^ in vertebrates [12, 15]. The corresponding six orders of magnitude variation in the drift barrier 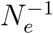 has immense ramifications for how we understand selection in prokaryotes versus eukaryotic organisms, particularly in the context of genome complexity [16–18]. For example, consider a mutant with an extra genetic sequence relative to the wild-type. We can separate *s* into two contributions, *s* = *s*_*c*_ + *s*_*a*_ [12]: *s*_*c*_ is the baseline selection coefficient associated with the metabolic costs of having this sequence, i.e. the costs of replicating it during cell division, synthesizing any associated mRNA / proteins, as well as the maintenance costs associated with turnover of those components; *s*_*a*_ is the correction due to any adaptive consequences of the sequence beyond its baseline metabolic costs. For a prokaryote with a low drift barrier 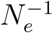, even the relatively low costs associated with replication and transcription are often under selective pressure [11, 12], unless *s*_*c*_ *<* 0 is compensated for an *s*_*a*_ *>* 0 of comparable or larger magnitude [19]. For the much greater costs of translation, the impact on growth rates of unnecessary protein production is large enough to be directly seen in experiments on bacteria [1, 20]. In contrast, for a eukaryote with sufficiently high 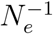, the same *s*_*c*_ might be effectively invisible to selection, even if *s*_*a*_ = 0. Thus even genetic material that initially provides no adaptive advantage can be readily fixed in a population, making eukaryotes susceptible to non-coding “bloat” in the genome. But this also provides a rich palette of genetic materials from which the complex variety of eukaryotic regulatory mechanisms can subsequently evolve [12, 21].

Part of the explanatory power of this idea is the fact that the *s*_*c*_ of a particular genetic variant should in principle be predictable from underlying physical principles. In fact, a very plausible hypothesis is that *s*_*c*_ *≈ -δC*_*T*_ */C*_*T*_, where *C*_*T*_ is the total resting metabolic expenditure of an organism per generation time, and *δC*_*T*_ is the extra expenditure of the mutant versus the wild-type. This relation can be traced at least as far back as the famous “selfish DNA” paper of Orgel and Crick [10], where it was mentioned in passing. But its true usefulness was only shown more recently, in the notable works of Wagner [11] on yeast and Lynch & Marinov [12] on a variety of prokaryotes and unicellular eukaryotes. By doing a detailed biochemical accounting of energy expenditures, they used the relation to derive values of *s*_*c*_ that provided intuitive explanations of the different selective pressures faced by different classes of organisms. The relation provides a Rosetta stone, translating metabolic costs into evolutionary terms. And its full potential is still being explored, most recently in describing the energetics of viral infection [22].

Despite its plausibility and long pedigree, to our knowledge this relation has never been justified in complete generality from first principles. We do so through a general bioenergetic growth model, versions of which have been applied across the spectrum of life [23–25], from unicellular organisms to complex vertebrates. We show that the relation is universal to an excellent approximation across the entire biological parameter range.

## Growth model

Let Π(*m*(*t*)) [unit: W] be the average power input into the resting metabolism of an organism (the metabolic expenditure after locomotion and other activities are accounted for [24]). Π(*m*(*t*)) can be an arbitrary function of the organism’s current mass *m*(*t*) [unit: g] at time *t*. This power is partitioned into maintenance of existing biological mass (i.e. the turnover energy costs associated with the constant replacement of cellular components lost to degradation), and growth of new mass (i.e. synthesis of additional components during cellular replication) [26]. Energy conservation implies

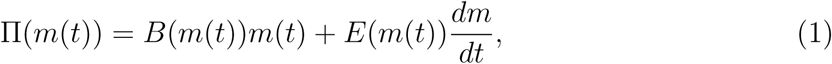

Here *B*(*m*(*t*)) [unit: W/g] is the maintenance cost per unit mass, and *E*(*m*(*t*)) [unit: J/g] is the synthesis cost per unit mass. We allow both these quantities to be arbitrary functions of *m*(*t*).

Though we will derive our main result for the fully general model of Eq. (1), we will also explore a special case: Π(*m*(*t*)) = Π_0_*m*^*α*^(*t*), *B*(*m*(*t*)) = *B*_*m*_, *E*(*m*(*t*)) = *E*_*m*_, with scaling exponent *α* and constants Π_0_, *B*_*m*_, and *E*_*m*_ [25]. Allometric scaling of Π(*m*(*t*)) with *α* = 3*/*4 across many different species was first noted in the work of Max Kleiber in the 1930s [27], and with the assumption of time-independent *B*(*m*(*t*)) and *E*(*m*(*t*)) leads to a successful description of the growth curves of many higher animals [23, 24]. However, recently there has been evidence that *α* = 3*/*4 may not be universal [28, 29]. Higher animals still exhibit *α <* 1 (with debate over *α* = 2*/*3 versus 3/4 [30]), but unicellular organisms have a broader range *α* ≲ 2. Thus we will use the model of Ref. [25] with an arbitrary species-dependent exponent *α*. While the resulting description is reasonable as a first approximation, particularly for unicellular organisms, one can easily imagine scenarios where the exponent and maintenance costs might vary between different developmental stages [31]. For the case of maintenance in endothermic animals, which in our approach includes all non-growth-related expenditures, more energy per unit mass is allocated to heat production as the organism matures [32], effectively increasing the cost of maintenance. In the Supplementary Information (SI) Sec. V [33] we show how the generalized model works in this scenario, using experimental growth data from two endothermic bird species [34]. Thus it is useful to initially consider the model in complete generality.

### Baseline selection coefficient for metabolic costs

To derive an expression for *s*_*c*_ for the growth model of Eq. (1), we first focus on the generation time *t*_*r*_, since this will be affected by alterations in metabolic costs. *t*_*r*_ is the typical age of reproduction, defined explicitly for any population model in SI Sec. I, where we relate it to the population birth rate *r* through *r* = ln(*R*_*b*_)*/t*_*r*_ [35, 36]. Here *R*_*b*_ is the mean number of offspring per individual. Let *ϵ* = *m*_*r*_*/m*_0_ be the ratio of the mass *m*_*r*_ = *m*(*t*_*r*_) at reproductive maturity to the birth mass *m*_0_ = *m*(0). For example in the case of symmetric binary fission of a unicellular organism, *R*_*b*_ *≈ ϵ ≈* 2 (see SI Sec. III for a discussion of *ϵ* in more general models of cell size homeostasis). Since *m*(*t*) is a monotonically increasing function of *t* for any physically realistic growth model, we can invert Eq. (1) to write the infinitesimal time interval *dt* associated with an infinitesimal increase of mass *dm* as *dt* = *dm E*(*m*)*/G*(*m*) where *G*(*m*) *≡* Π(*m*) *- B*(*m*)*m* is the amount of power channeled to growth, and we have switched variables from *t* to *m*. Note that *G*(*m*) must be positive over the *m* range to ensure that *dm/dt >* 0. Integrating *dt* gives us an expression for *t*_*r*_,

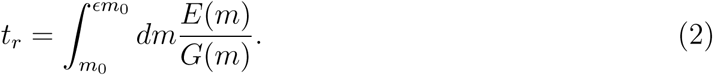

If we are interested in finding *s*_*c*_ for a genetic variation, we can focus on the additional metabolic costs due to that variation. For the purposes of calculation, this means treating the mutation as if it does not alter biological function in any other respect, including the ability of the organism to assimilate energy for its resting metabolism through uptake of nutrients or foraging. If the mutation actually had only metabolic cost effects, the full selection coefficient *s* = *s*_*c*_. However generically mutations can affect both metabolic costs and power input (and/or other adaptive aspects), so *s* = *s*_*c*_ + *s*_*a*_, with a correction term *s*_*a*_ due to the adaptive effects [12]. In the latter case *s*_*c*_ can still be calculated as shown below (ignoring adaptive effects) and interpreted as the baseline contribution to selection due to metabolic costs. While we do not focus on *s*_*a*_ here, our theory can be readily extended to consider adaptive contributions as well, as illustrated in SI Sec. VII, including aspects like spare respiratory capacity. This broader formalism is summarized in Fig. S3.

Proceeding with the *s*_*c*_ derivation, the products of the genetic variation (i.e. extra mRNA transcripts or translated proteins) may alter the mass of the mutant, which we denote by 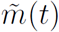. The left-hand side of Eq. (1) remains Π(*m*(*t*)), where *m*(*t*) is now the *unperturbed* mass of the organism (the mass of all the pre-variation biological materials). The power input Π(*m*(*t*)) depends on *m*(*t*) rather than 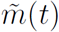 since only *m*(*t*) contributes to the processes that allow the organism to process nutrients, in accordance with the assumption that power input is unaltered in order to calculate *s*_*c*_. It is also convenient to express our dynamics in terms of *m*(*t*) rather than 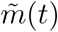, since the condition defining reproductive time *t*_*r*_ remains unchanged, *m*(*t*_*r*_) = *Em*_0_, or in other words when the unperturbed mass reaches *ϵ* times the initial unperturbed mass *m*_0_. Thus Eq. (1) for the mutant takes the form 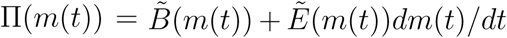, where 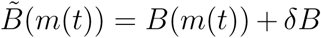 and 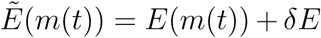 are the mutant maintenance and synthesis costs. For simplicity, we assume the perturbations *δB* and *δE* are independent of *m*(*t*), though this assumption can be relaxed. In SI Sec. IV, we show a sample calculation of *δB* and *δE* for mutations in *E. coli* and fission yeast involving short extra genetic sequences transcribed into non-coding RNA. This provides a concrete illustration of the framework we now develop.

Changes in the metabolic terms will perturb the generation time, 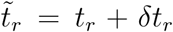, and consequently the birth rate 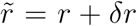. The corresponding baseline selection coefficient *s*_*c*_ can be exactly related to 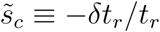, the fractional change in *t*_*r*_, through 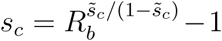 (see SI Sec. I). This relation can be approximated as *s*_*c*_ *≈* ln 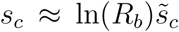 when 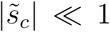, the regime of interest when making comparisons to drift barriers 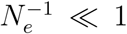. In this regime 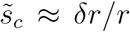, the fractional change in birth rate. While we focus here on the the simplest case of exponential population growth, where 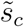 is time-independent, we generalize our approach to density-dependent growth models, where 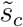 varies between generations, in SI Sec. VI. 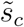 can be written in a way that directly highlights the contributions of *δE* and *δB* to 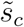. To facilitate this, let us define the average of any function *F* (*m*(*t*)) over a single generation time *t* as 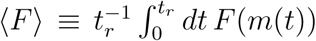. Changing variables from *t* to *m*, like we did above in deriving Eq. (2), we can write this equivalently as 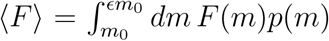, where 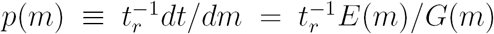. The value *p*(*m*)*dm* is just the fraction of the generation time that the organism spends growing from mass *m* to mass *m* + *dm*. Expanding Eq. (2) for *t*_*r*_ to first order in the perturbations *δE* and *δB*, the coefficient 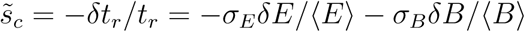, with positive dimensionless prefactors

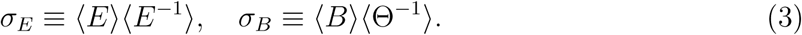

Here Θ(*m*) *≡ G*(*m*)*/m*, and *F* ^*-*1^(*m*) *≡* 1*/F* (*m*) for any *F*. The magnitude of *σ*_*B*_ versus *σ*_*E*_ describes how much fractional increases in maintenance costs matter for selection relative to fractional increases in synthesis costs. We see that both prefactors are products of time averages of functions related to metabolism. See SI Sec. II for a detailed derivation of Eq. (3), and also Eq. (4) below.

## Relating the baseline selection coefficient to the fractional change in total resting metabolic costs

The final step in our theoretical framework is to connect the above considerations to the total resting metabolic expenditure *C*_*T*_ of the organism per generation time *t*_*r*_, given by 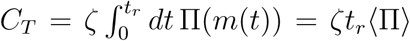. To compare with the experimental data of Ref. [12], compiled in terms of phosphate bonds hydrolyzed [P], we add the prefactor *ζ* which converts from units of J to P. Assuming an ATP hydrolysis energy of 50 kJ/mol under typical cellular conditions, we set *ζ* = 1.2 *×* 10^19^ P/J. The genetic variation discussed above perturbs the total cost, 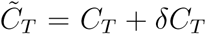, and the fractional change *δC*_*T*_ */C*_*T*_ can be expressed in a form analogous to 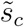, namely *δC*_*T*_ */C*_*T*_ = *σE’ E/*⟨*E*⟩ + *σ ′* _*B*_ *δB/*⟨*B*⟩, with

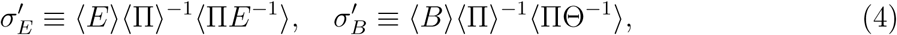

where again the prefactors are expressed in terms of time averages over metabolic functions. The connection between *s*_*c*_ and *δC*_*T*_ */C*_*T*_ can be constructed by comparing Eq. (3) with Eq. (4). We see that 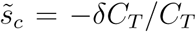 for all possible perturbations *δE* and *δB* only when *σ*_*E*_ = *σ′*_*E*_ and *σ*_*B*_ = *σ′*_*B*_ We derive strict bounds on the differences between the prefactors (SI Sec. II), which show that the relation is exact when: i) Π(*m*) is a constant independent of *m*; and/or ii) *E*(*m*) and Θ(*m*) are independent of *m*. Outside these cases, the relation 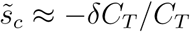 is an approximation. To see how well it holds, it is instructive to investigate the allometric growth model described earlier, where Π(*m*(*t*)) = Π_0_*m*^*α*^(*t*), *E*(*m*(*t*)) = *E*_*m*_, *B*(*m*(*t*)) = *B*_*m*_.

## Testing the relation in an allometric growth model

We use model parameters based on the metabolic data of Ref. [12], covering a variety of prokaryotes and unicellular eukaryotes. This data consisted of two quantities, *C*_*G*_ and *C*_*M*_, which reflect the growth and maintenance contributions to *C*_*T*_. Using Eq. (1) to decompose Π(*m*(*t*)), we can write *C*_*T*_ = *C*_*G*_+ *t*_*r*_ *C*_*M*_, where 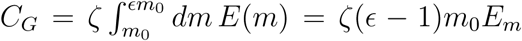 is the expenditure for growing the organism, and *C*_*M*_ = *ζ* ⟨ *Bm* ⟩ = *ζB*_*m*_ ⟨*m* ⟩ is the mean metabolic expenditure for maintenance per unit time. *C*_*G*_ and *C*_*M*_ scale linearly with cell volume (SI Sec. III), and best fits to the data, shown in Fig. 1, yield global interspecies averages: *E*_*m*_ = 2, 600 J/g and *B*_*m*_ = 7 *×* 10^*-*3^ W/g. As discussed in the SI, these values are remarkably consistent with earlier, independent estimates, for unicellular and higher organisms [24, 25, 37, 38].

**FIG. 1.**
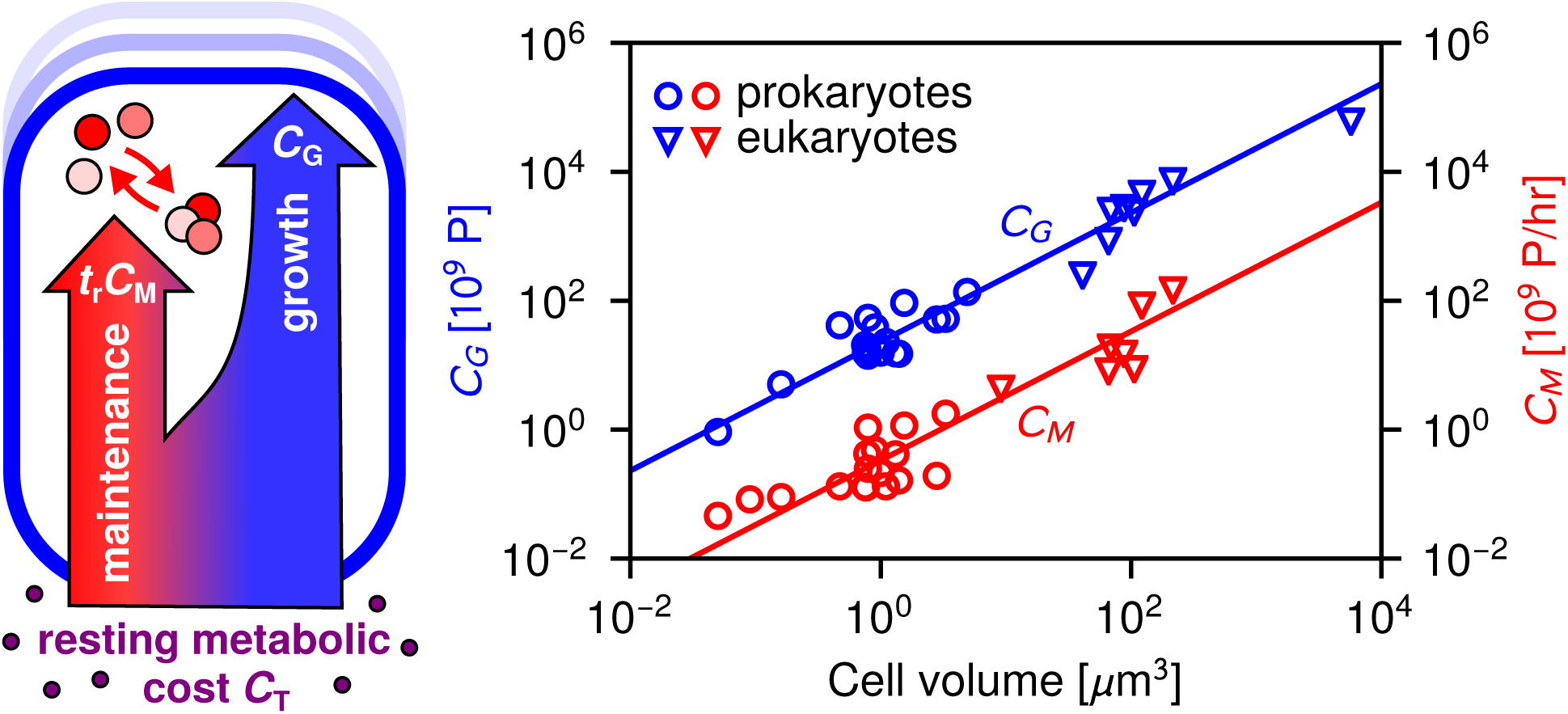
The growth *C*_*G*_ (blue) and maintenance *C*_*M*_ (red) contributions to an organism’s total resting metabolic cost *C*_*T*_ = *C*_*G*_ +*t*_*r*_*C*_*M*_ per generation time *t*_*r*_. The symbols (circles = prokaryotes, triangles = unicellular eukaryotes) represent data tabulated in Ref. [12]. *C*_*G*_ and *C*_*M*_ have units of 10^9^ P (phosphate bonds hydrolyzed), and 10^9^ P/hr respectively. The lines represent best fits to the theoretical expressions for *C*_*G*_ and *C*_*M*_ from the allometric growth model.

Since *E*(*m*(*t*)) = *E*_*m*_ is a constant in the allometric growth model, *σ*_*E*_ = 1 from Eq. (3), and *σ*_*E*_ = *σ′E* holds exactly from Eq. (4). So the only aspect of the approximation that needs to be tested is the similarity between *σ*_*B*_ and *σ′B*. Fig. 2A shows *σ*_*B*_ versus *σ′B* for the range *α* = 0 *-* 3, which includes the whole spectrum of biological scaling [28] up to *α* = 2, plus some larger *α* for illustration. For a given *α*, the coefficient Π_0_ has been set to yield a certain division time *t*_*r*_ = 1 *-* 40 hr, encompassing both the fast and slow extremes of typical unicellular reproductive times. In all cases *σ′B* is in excellent agreement with *σ*_*B*_. For the range *α ≤* 2 the discrepancy is less than 15%, and it is in fact zero at the special points *α* = 0, 1. Clearly the approximation begins to break down at *α "* 1, but it remains sound in the biologically relevant regimes. Note that *σ*_*B*_ values for *t*_*r*_ = 1 hr are *∼* 0.01, reflecting the minimal contribution of maintenance relative to synthesis costs in determining the selection coefficient for fast-dividing organisms. This limit is consistent with microbial metabolic flux theory [39], where maintenance is typically neglected, so 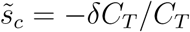 exactly (since only *σ*_*E*_ = *σ*_*E*_′= 1 matters). As *t*_*r*_ increases, so does *σ*_*B*_ and hence the influence of maintenance costs, so by *t*_*r*_ = 40 hr, *σ*_*B*_ is comparable to *σ*_*E*_.

**FIG. 2.**
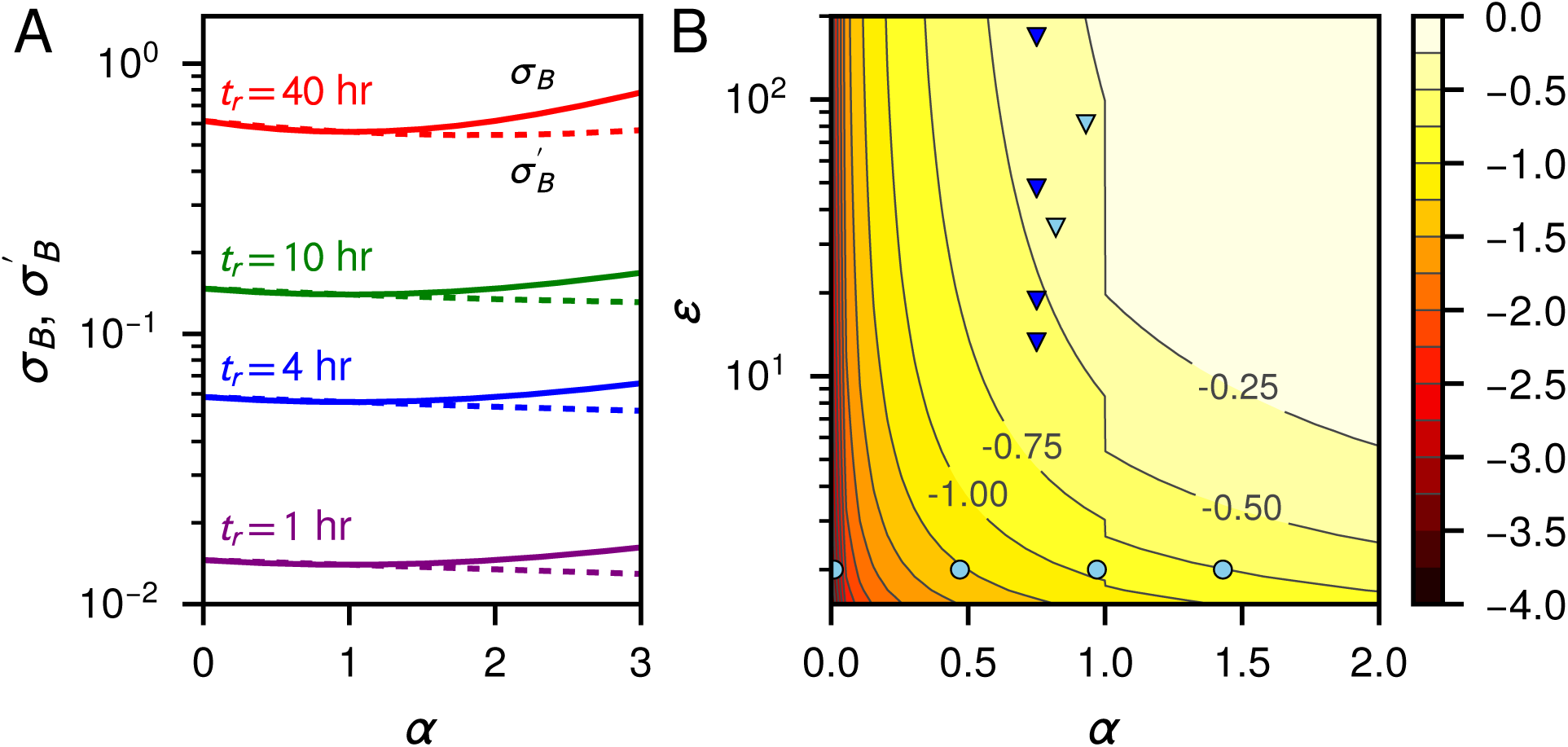
A: *σ*_*B*_ (solid curves) from Eq. (3) and *σ′B* (dashed curves) from Eq. (4) versus *α*, for the allometric growth model with *E*_*m*_ = 2, 600 J/g, *B*_*m*_ = 7 *×* 10^*-*3^ W/g, and ϵ = 2. At any given *α*, the parameter Π_0_ for each pair of curves (different colors) is chosen to correspond to particular reproductive times *t*_*r*_, indicated in the labels. B: Contour diagram showing the logarithm of the maximum possible discrepancy log10 *|*1 *- σ′B /σ*_*B*_*|* for any allometric growth model parameters, as a function of *α* and *E*. To illustrate biological ranges *α* and *E*, the symbols correspond to data for various species (circles = unicellular, triangles = multicellular) drawn from the growth trajectories analyzed in Ref. [25] (light blue) and Ref. [23] (dark blue). SI Sec. III shows a detailed species list.

To make a more comprehensive analysis of the validity of the 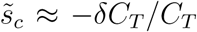 relation, we do a computational search for the worst case scenarios: for each value of *α* and *E*, we can numerically determine the set of other growth model parameters that gives the largest discrepancy *|*1 *- σ′*_*B*_ */σ*_*B*_*|*. Fig. 2B shows a contour diagram of the results on a logarithmic scale, log10 *|*1 *- σ′*_*B*_ */σ*_*B*_*|*, as a function of *α* and *E*. Estimated values for *α* and *E* from the growth trajectories of various species are plotted as symbols to show the typical biological regimes. While the maximum discrepancies are smaller for the parameter ranges of unicellular organisms (circles) compared to multicellular ones (triangles), in all cases the discrepancy is less than 50%. To observe a serious error (*σ′*_*B*_ a different order of magnitude than *σ*_*B*_), one must go to the large *α*, large *E* limit (top right of the diagram) which no longer corresponds to biologically relevant growth trajectories.

## Validity of the relation in more complex growth scenarios

Going beyond the simple allometric model, SI Sec. V analyzes avian growth data, where the metabolic scaling exponent varies between developmental stages. We find *σ*_*E*_ = *σ′*_*E*_ = 1 and the discrepancy *|*1 *- σ′*_*B*_ */σ*_*B*_*| ≤* 30%. SI Sec. VI considers density-dependent growth, illustrated by examples of bacteria competing for a limited resource in a chemostat and predators competing for prey. Remarkably, when these systems approach a stationary state in total population and resource/prey quantity, we find *σ*_*E*_ = *σ′*_*E*_ = 1, *σ*_*B*_ = *σ′*_*B*_ = (*B*_*m*_ ln *R*_*b*_)*/*(*E*_*m*_*d* ln ϵ), where *d* is the dilution rate in the chemostat, or the predator death rate. The simple expression for *σ*_*B*_ allows straightforward estimation of the maintenance contribution to selection. For the chemostat that contribution can be tuned experimentally through the dilution rate *d*.

## Conclusion

We thus reach the conclusion that the baseline selection coefficient for metabolic costs can be reliably approximated as *s*_*c*_ *≈ -* ln(*R*_*b*_)*δC*_*T*_ */C*_*T*_. As in the original hypothesis [10–12], *-δC*_*T*_ */C*_*T*_ is the dominant contribution to the scale of *s*_*c*_, with corrections provided by the logarithmic factor ln(*R*_*b*_). Our derivation puts the relation for *s*_*c*_ on a solid footing, setting the stage for its wider deployment. It deserves a far greater scope of applications beyond the pioneering studies of Refs. [11, 12, 22]. Knowledge of *s*_*c*_ can also be used to deduce the adaptive contribution *s*_*a*_ = *s - s*_*c*_ of a mutation, which has its own complex connection to metabolism [40] (see also SI Sec. VII). The latter requires measurement of the overall selection coefficient *s*, for example from competition/growth assays, and the calculation of *s*_*c*_ from the relation, assuming the underlying energy expenditures are well characterized. The *s*_*c*_ relation underscores the key role of thermodynamic costs in shaping the interplay between natural selection and genetic drift. Indeed, it gives impetus to a major goal for future research: a comprehensive account of those costs for every aspect of biological function, and how they vary between species, what one might call the “thermodynome”. Relative to its more mature omics brethren—the genome, proteome, transcriptome, and so on—the thermodynome is still in its infancy, but fully understanding the course of evolutionary history will be impossible without it.

The authors thank useful correspondence with M. Lynch, and feedback from B. Kuznets-Speck, C. Weisenberger, and R. Snyder. E.I. acknowledges support from Institut Curie.

## Supplementary Information

### I. Derivation of the relation between *s*_*c*_ and 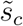

In the main text we posited an approximate relation 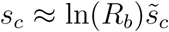 between the baseline selection coefficient *s*_*c*_ and the fractional change in growth rate 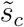 due to a genetic variation. Here we derive an exact relation between the two quantities, generalizing the approach used in Ref. [77] for the specific case of binary fission. We then show how the approximation used in the main text arises in the limit 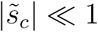.

Consider a group of wild-type organisms with population *N*_*w*_(*t*) as a function of time. Over each generation time *t*_*r*_ the population grows by a factor of *R*_0_, the net reproductive rate, so that after *n* generations,

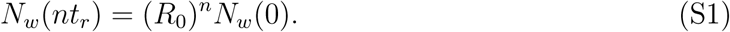

Here we focus on the simple case of exponential growth where both *R*_0_ and *t*_*r*_ are time-independent. Below in Sec. VI we consider more complex scenarios in systems with density-dependent population growth. For a growing population *R*_0_ *>* 1. Most generally *R*_0_ and *t*_*r*_ are defined through 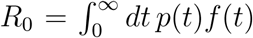 and 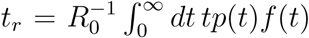, where *p*(*t*) is the probability of the organism to survive from birth to age *t*, and *f* (*t*) is the average fecundity at age *t* [35, 36]. A common simplification is to assume *f* (*t*) is sharply peaked around age *t*_*r*_, *f* (*t*) *≈ R*_*b*_*δ*(*t - t*_*r*_) with mean number of offspring *R*_*b*_, so that *R*_0_ *≈ R*_*b*_*p*(*t*_*r*_) [25, 36]. One example is binary fission in bacteria, where *R*_*b*_ = 2, and *p*(*t*_*r*_) *≈* 1 if one neglects cell deaths (and there is no cell removal), so *R*_0_ *≈ R*_*b*_ = 2. In the continuum time approximation to population growth, *n* = *t/t*_*r*_ and Eq. (S1) becomes

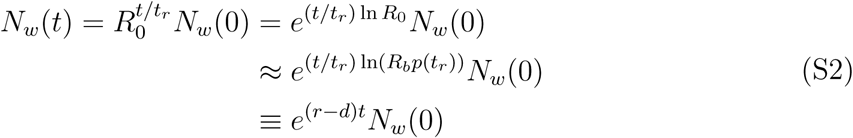

describing Malthusian growth with net rate *r - d*, the difference between the birth rate *r* = ln(*R*_*b*_)*/t*_*r*_ and death rate *d* = *-* ln(*p*(*t*_*r*_))*/t*_*r*_.

Now consider the growth of a mutant population *N*_*m*_(*t*). Under the baseline assumption on which we focus in the main text (neglecting any adaptive effects), the mutant has the same mean number of offspring *R*_*b*_ and death rate *d*, but metabolic cost differences affecting growth perturb the mean generation time to 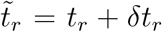. This leads to a modified birth rate 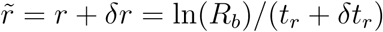 ln(*R*_*b*_)*/*(*t*_*r*_ + *δt*_*r*_), and a growth equation

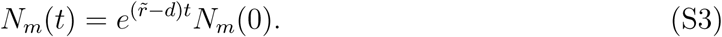

From Eqs. (S2)-(S3) and the definitions of *r* and 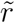, the ratio of the mutant to wild type populations is given by

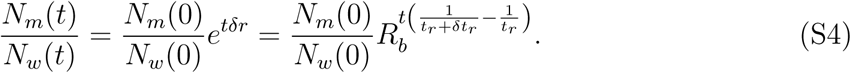

On the other hand, after *n* wild-type generations (*t* = *nt*_*r*_) the ratio of the two populations is related to the selection coefficient (as conventionally defined in population genetics) through

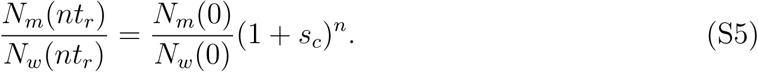

Plugging *t* = *nt*_*r*_ into Eq. (S4) and comparing to Eq. (S5), we see that

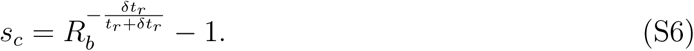

If we define 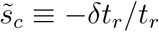, then Eq. (S6) can be rewritten as

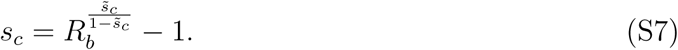

Note that in the case where 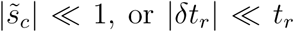, we can also write 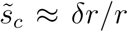, and so interpret 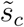 as the fractional change in birth rate. In this same limit we can expand Eq. (S7) for small 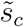,

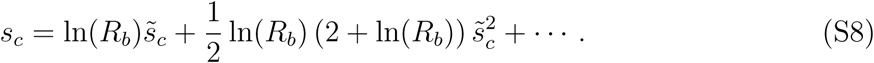

Keeping only the leading order term, linear in 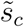, yields the approximation *s*_*c*_ *≈* ln 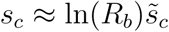.

### II. Derivation of main text Eqs. (3)-(4) and related bounds

To derive Eq. (3) in the main text, we start with the equation for *t*_*r*_ [main text Eq. (2)]:

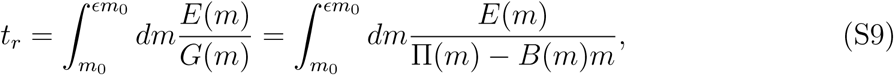

with *G*(*m*) *≡* Π(*m*) *- B*(*m*)*m*. Under the perturbations *E*(*m*) *→ E*(*m*) + *δE* and *B*(*m*) *→ B*(*m*) + *δB*, the generation time is altered to

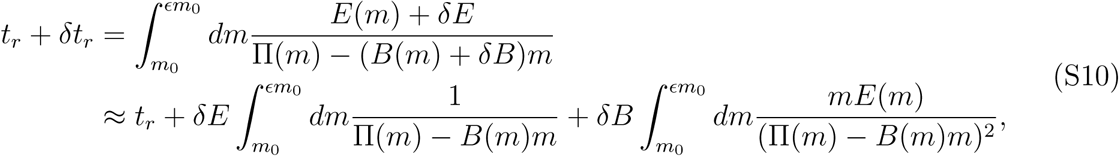

where in the second line we have Taylor expanded the integrand to first order in *δE* and *δB*. We can thus write 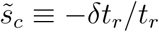 as

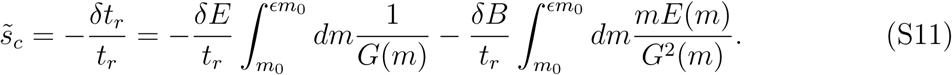

Using the definitions 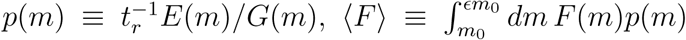, and Θ(*m*) *≡ G*(*m*)*/m*, we can rewrite Eq. (S11) as

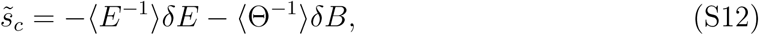

where *F* ^*-*1^(*m*) *≡* 1*/F* (*m*) for any function *F*. Finally we define the prefactors *σ*_*E*_ and *σ*_*B*_ through the relation

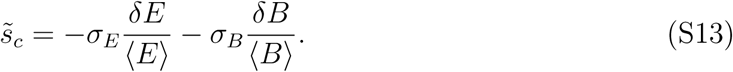

Comparing Eq. (S12) to (S13) we find the result of main text Eq. (3):

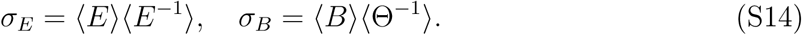

The derivation of main text Eq. (4) proceeds analogously, starting with the total resting metabolic expenditure per generation,

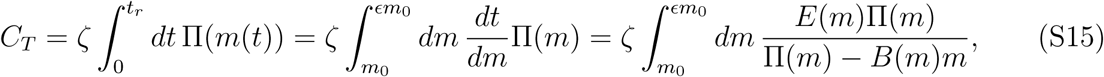

where we have changed variables in the integral from *t* to *m* and used the fact that *dt/dm* = *E*(*m*)*/G*(*m*). Under the perturbations *E*(*m*) *→ E*(*m*) + *δE* and *B*(*m*) *→ B*(*m*) + *δB*, the expenditure is altered to

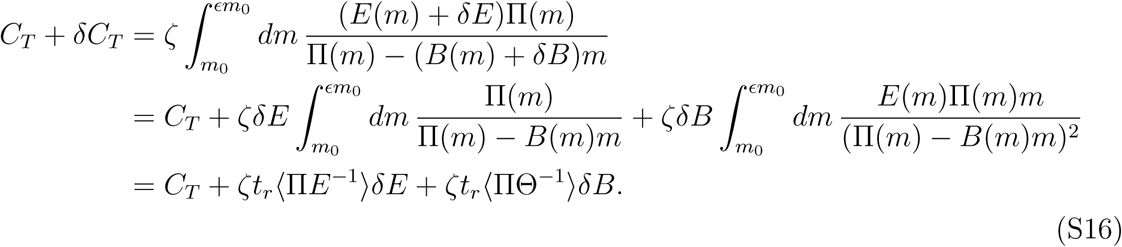

Using the fact that Eq. (S15) can also be written as *C*_*T*_ = *ζt*_*r*_⟨Π⟩, we can use Eq. (S16) to express the ratio *δC*_*T*_ */C*_*T*_ as

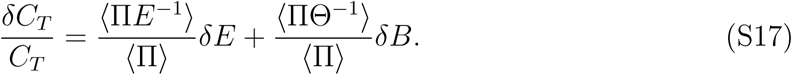

Comparing this to the expression defining *σ′*_*E*_ and *σ′*_*B*_,

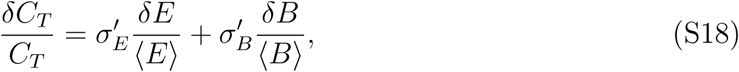

we find the result of main text Eq. (4):

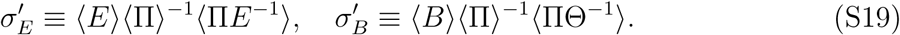

The degree to which 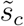 can be approximated as *δC*_*T*_ */C*_*T*_ depends on the similarity of the prefactor *σ*_*E*_ to *σ′*_*E*_, and *σ*_*B*_ to *σ′*_*B*_. Their relative differences can be written as:

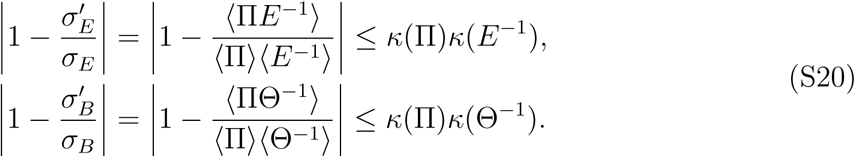

where 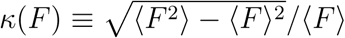 and we have used the Cauchy-Schwarz inequality. These bounds imply two cases when 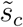 is exactly equal to *δC*_*T*_ */C*_*T*_ : i) *κ*(Π) = 0, which means Π(*m*) is a constant independent of *m*; ii) *κ*(Π) *>* 0 and *κ*(*E*^*-*1^) = *κ*(Θ^*-*1^) = 0, which means *E*(*m*) and Θ(*m*) are independent of *m*.

### III. Fitting of allometric growth model to experimental data

As discussed in the main text, we can decompose *C*_*T*_ into two components, *C*_*T*_ = *C*_*G*_ + *t*_*r*_*C*_*M*_, where 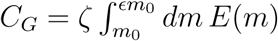 is the expenditure for growing the organism, and *C*_*M*_ = *ζ*⟨*Bm*⟩ is the mean metabolic expenditure for maintenance per unit time. For the allometric growth model, these contributions to *C*_*T*_ simplify to *C*_*G*_ = *ζ*(ϵ *-*1)*m*_0_*E*_*m*_ and *C*_*M*_ = *ζB*_*m*_⟨*m*⟩. Ref. [12] noted that *C*_*G*_ and *C*_*M*_ collected from experimental data scaled nearly linearly with cell volume, with allometric exponents of 0.97 *±* 0.04 and 0.88 *±* 0.07 respectively. In fact, the simplest version of the allometric model predicts exactly linear scaling, using the following assumptions. Since the data tabulated in Ref. [12] covers prokaryotes and unicellular eukaryotes, we take ϵ = 2. Since the mass of the organism varies between *m*_0_ and 2*m*_0_ over time *t*_*r*_, we approximate ⟨*m*⟩ *≈* (3*/*2)*m*_0_. Note that setting ϵ = 2 assumes symmetric binary fission, though unicellular organisms can also exhibit asymmetric fission where ϵ *≠* 2 [57]. However since the variation in *E* is typically within a factor of two of the symmetric case, any errors introduced by this assumption, and the approximation for ⟨*m*⟩, will not change the order of magnitude of the estimated model parameters. For simplicity we are also ignoring variation in birth sizes *m*_0_ over the population, which in the unicellular case is closely related to the possible mechanisms of cell size homeostasis, so-called sizer, adder, or timer behaviors [47, 48]. In the current context, once a steady state size distribution is reached, we can define *E* as the ratio of mean size at division to the mean birth size *m*_0_, which will have different interpretations depending on the mechanism. In the sizer picture ϵ = *m*_r_*/m*_0_, where *m*_r_ is the target cell mass at which division occurs. In the adder picture, where cells divide after adding a mass Δ*m*, we have *ϵ* = 1+Δ*m/m*_0_. In the timer mechanism, where division occurs after a specified time, additional conditions are needed to achieve a steady state size distribution, for example linear growth (in which case timer and adder are equivalent) or a mixed model coupling sizer and timer behavior [41]. In all cases, capturing the full effects of these mechanisms would require modeling the variation in birth sizes and their evolution over time, which we leave for a future work.

We relate the experimentally observed cell volume *V* to the mean cell mass ⟨*m*⟩ by assuming a typical cell is 2/3 water (density *ρ*_wat_ = 10^*-*12^ g*/µ*m^3^) and 1/3 dry biomass (density *ρ*_dry_ *≈* 1.3 *×* 10^*-*12^ g*/µ*m^3^) [80]. Hence ⟨*m*⟩ = (2*ρ*_wat_ + *ρ*_dry_)*V/*3 *≡ ρ*_cell_*V*. We thus find:

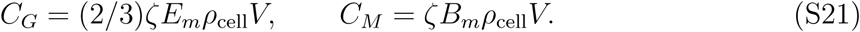

For each expression we have only one unknown parameter, *E*_*m*_ and *B*_*m*_ respectively. Best fits to the Ref. [12] data, shown in main text Fig. 1B, yield global interspecies averages of the parameters, *E*_*m*_ = 2, 600 J/g and *B*_*m*_ = 7 *×* 10^*-*3^ W/g.

The fitted values are consistent with earlier approaches, once water content is accounted for (i.e. to get *E*_*m*_ per dry biomass, multiply the value by *≈* 3, so 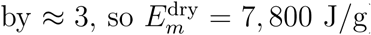). The synthesis cost *E*_*m*_ has a very narrow range across many species, with *E*_*m*_ = 1, 100 *-* 1, 800 J/g in bird and fish embryos, and 4, 000 *-* 7, 500 J/g for mammal embryos and juvenile fish, birds, and mammals [37]. This energy scale seems to persist down to the prokaryotic level, with 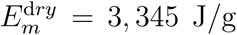 J/g estimated for *E. coli* [25]. 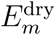 also appears in a different guise as the inverse of the “energy efficiency” *ε* of *E. coli* growth in the model of Ref. [38]; converting the optimal observed *ε ≈* 15 dry g/(mol ATP) yields 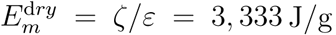 J/g, consistent with the other estimates cited above, as well as our fitted value. The ratio *B*_*m*_*/E*_*m*_ was estimated for various species in Ref. [25], and found to vary in the range 10^*-*6^ *-* 10^*-*5^ s^*-*1^ from prokaryotes to unicellular eukaryotes, entirely consistent with our fitted value of *B*_*m*_*/E*_*m*_ = 3 *×* 10^*-*6^ s^*-*1^. The scale shifts for larger, multicellular species, but not dramatically. For example for a subset of mammals with scaling *α* = 3*/*4, adult mass sizes *m*_*a*_ = 10 *-*6.5 *×*10^5^ g, and typical values of Π_0_ *≈* 0.022 W/g^3*/*4^, *E*_*m*_ *≈* 7000 J/g [24], we get a range of *B*_*m*_*/E*_*m*_ = 10^*-*7^ *-* 10^*-*6^ s^*-*1^. We thus have confidence that the growth model provides a description of the metabolic expenditures (in terms of growth and maintenance contributions) that is consistent both with the empirical data of Ref. [12] and parameter expectations based on a variety of earlier approaches.

For the symbols in the contour diagram of main text figure Fig. 2B, we used parameters extracted from growth trajectories analyzed in Ref. [25] (light blue) and Ref. [23] (dark blue). Circles (left to right) are unicellular organisms (*ϵ* = 2): *T. weissflogii, L. borealis, B. subtilis, E. coli*. Triangles (top to bottom) are multicellular organisms: guinea pig, *C. pacificus*, hen, *Pseudocalanus sp.*, guppy, cow. For the multicellular case the plotted values of *ϵ* correspond to asymptotic adult mass in units of *m*_0_. This is an upper bound on ϵ, though the actual *ϵ* should typically be comparable [23, 78].

### IV. Sample calculation of the baseline selection coefficient: short, non-coding RNA in *E. coli* **and fission yeast**

To illustrate a calculation of baseline selection coefficients in the framework developed in the main text, let us consider a specific biological example: a mutant with a short (*<* 200 bp) sequence in the genome that is transcribed into non-coding RNA, and which is not present in the wild-type. We will focus on two organisms, the prokaryote *E. coli* and the unicellular eukaryote *S. pombe* (fission yeast). To date we know that at least some subset of non-coding RNA transcripts have functional roles in these organisms [75, 76]. The evolution of such regulatory sequences will be shaped both by the selective advantage *s*_*a*_ of having the sequence in the genome, and the baseline disadvantage *s*_*c*_ from the extra energetic costs of copying and transcription.

Before calculating *s*_*c*_, we first establish the validity of the growth model for these organisms. The model parameters fitted for the data from prokaryotes and unicellular eukaryotes in Fig. 1B of the main text are *E*_*m*_ = 2, 600 J/g and *B*_*m*_ = 7 *×* 10^*-*3^ W/g. The corresponding growth and maintenance contributions to the total resting metabolic cost per generation, *C*_*G*_ and *C*_*M*_, are given by Eq. (S21). Using *ζ* = 1.2 *×* 10^19^ P/J (recall that P corresponds to ATP or ATP equivalents hydrolyzed), *ρ*_cell_ = 1.1 *×* 10^*-*12^ g/*µ*m^3^, and typical cell volumes *V*_*E.coli*_ = 1 *µ*m^3^ [80], *V*_*S.pombe*_ = 106 *µ*m^3^ [74], we find: 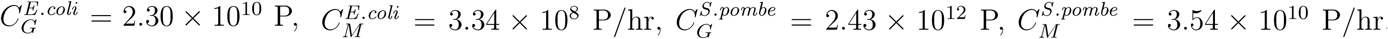. These agree well in magnitude with the literature estimates compiled in the SI of Ref. [12] (all normalized to 20°C): 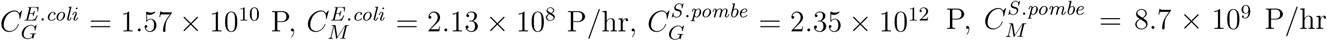. Thus the globally fitted *E*_*m*_ and *B*_*m*_ values are physically reasonable for both organisms.

The extra sequence in the mutant leads to perturbations in both synthesis cost per unit mass, *δE*, and maintenance cost per unit mass, *δB*. To calculate the first, we use the following estimates based on the analysis in Ref. [12]: for a sequence of length *L*, the total DNA-related synthesis cost is *d*_*ξ*_*L*, where the label *ξ* = *E. coli* or *S. pombe*. Here the prefactor *d*_*E.coli*_ *≈* 101 P and *d*_*S.pombe*_ *≈* 263 P. If the steady-state average number of corresponding mRNA trascripts in the cell is *N*_*r*_, the additional ribonucleotide synthesis costs are *≈* 46*N*_*r*_*L* in units of P. Hence we have, per unit mass,

**FIG. S1.**
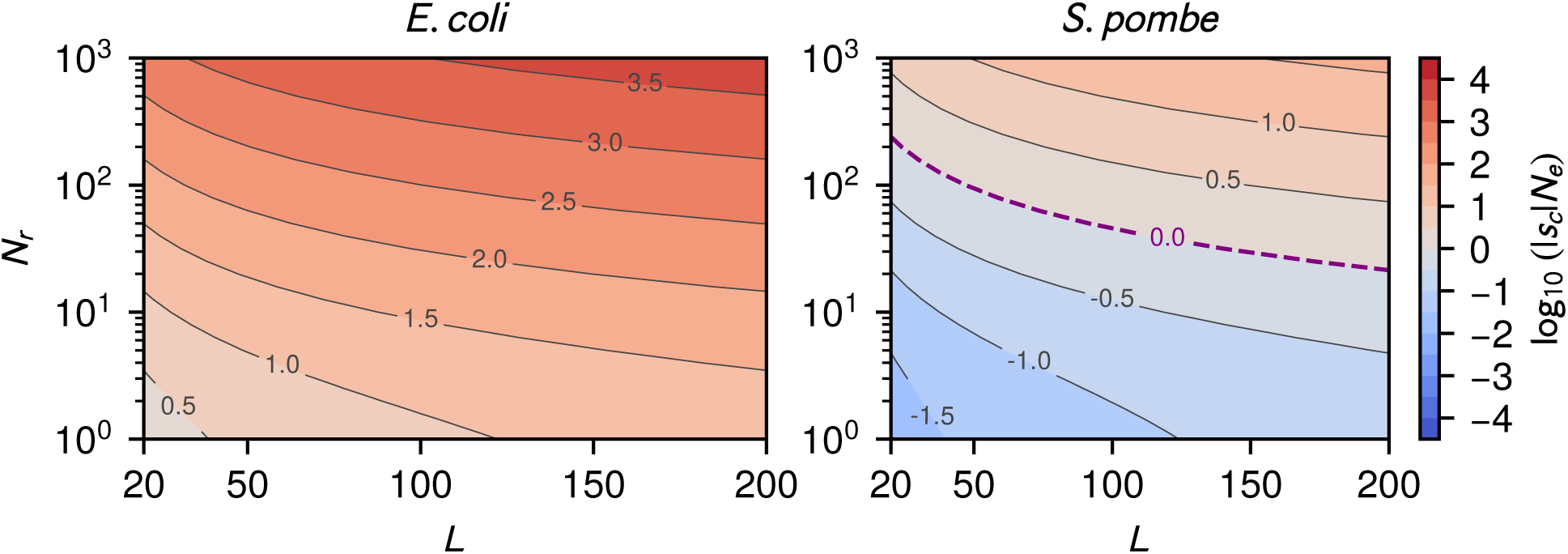
Contour diagrams of log_*10*_(*|s*_*c*_*|N*_*e*_) as a function of sequence length *L* and mean RNA transcript number *N*_*r*_ per cell for *E. coli* (left) and *S. pombe* (right). The dashed line in the diagram on the right corresponds to 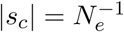.

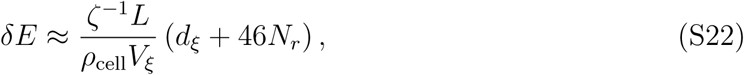

with the *ζ*^*-*1^ prefactor converting from P to J, so that *δE* has units of J/g. The same analysis [12] yields the maintenance cost per unit time for replacing transcripts after degradation, *≈* 2*N*_*r*_*Lγ*_*ξ*_ in units of P/s, where *γ*_*E.coli*_ = 0.003 s^*-*1^ and *γ*_*S.pombe*_ = 0.001 s^*-*1^ are the RNA degradation rates for the two organisms. Per unit mass, the maintenance perturbation *δB* is given by

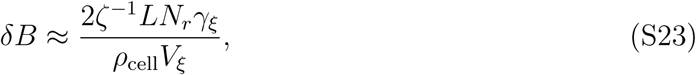

in units of W/g.

The final step is to calculate the prefactors *σ*_*E*_ and *σ*_*B*_ from Eq. (3) in the main text. For this we need to choose a particular growth model exponent *α*, and we set *α* = 1, corresponding to the assumption of exponential cell mass growth. In this case *σ*_*E*_ = 1 for both organisms, while 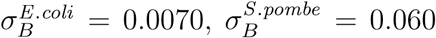.The choice of *α* has a minimal influence on the prefactors: *σ*_*E*_ = 1 exactly for any model with a constant function *E*(*m*) = *E*_*m*_. Moreover, any *α* value in the biologically relevant range of 0 *≤ α ≤* 2 yields a *σ*_*B*_ value within 5% of the *α* = 1 result for each organism.

Putting everything together, we now can calulate all the components of main text Eq. (3) for 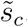, namely *σ*_*E*_, *σ*_*B*_, *δ*_*E*_, *δ*_*B*_, ⟨*E*⟩ = *E*_*m*_, and ⟨*B*⟩ = *B*_*m*_. Had we chosen instead to use the *δC*_*T*_ */C*_*T*_ approximation of main text Eq. (4), the only discrepancy would have been in the fact that *σ′*_*B*_ ≠ *σ*_*B*_, since *σ′*_*E*_ = *σ*_*E*_ = 1. However the discrepancy is small, with *|*1 - *σ*′_*B*_ /*σ*_*B*_*| <* 0.09 for both organisms in the range 0 *≤ α ≤* 2.

Fig. S1 shows contour diagrams of log_10_(|*s*_*c*_|*N*_*e*_) as a function of *L* and *N*_*r*_ for *E. coli* and *S. pombe*. Here 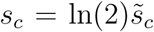, assuming *R*_*b*_ = 2, and the effective population sizes are 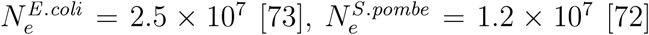. For *E.coli*, with its smaller metabolic expenditures per generation relative to fission yeast, the cost of the extra sequence is always significant: 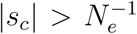 for the entire range of *L* and *N*_*r*_ considered, even for the smallest length (*L* = 20 bp) and a single transcript per cell on average, *N*_*r*_ = 1. Thus there will always be strong selective pressure to remove the extra sequence, unless *s*_*c*_ is compensated for by a comparable or greater adaptive advantage *s*_*a*_. In contrast, for *S. pombe* there is a regime of *L* and *N*_*r*_ where 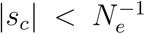 (the region below the dashed line). Here the selective disadvantage of the extra sequence is weaker than genetic drift, and such a genetic variant could fix in the population at roughly the same rate as a neutral mutation even if it conferred no selective advantage, *s*_*a*_ = 0. While this makes fission yeast more tolerant of genomic “bloat” relative to *E. coli*, initially non-functional extra genetic material could subsequently facilitate the development of novel regulatory mechanisms.

### V. Generalized growth model with varying developmental scaling regimes

In the last section of the main text, we explored the validity of the baseline selection coefficient relation in the simplest version of the growth model, with allometric power input Π(*m*(*t*)) = Π_0_*m*^*α*^(*t*) and time-independent synthesis and maintenance factors *E*(*m*(*t*)) = *E*_*m*_, *B*(*m*(*t*)) = *B*_*m*_. While this may be a reasonable approximation for various kinds of organisms [23–25] (particularly those without major physiological / morphological changes during their development [31]), there is evidence that in certain cases a single power law scaling with mass cannot accurately capture the resting metabolic rate (a detailed review is provided in Ref. [31]). This includes insects [69, 70] and marine invertebrates [71] that progress through several distinct developmental stages or instars, as well as endothermic birds when comparing juvenile and adult metabolism [34]. In these cases the resting metabolic input Π(*m*(*t*)) may scale approximately like a power-law in *m*(*t*) during individual stages of development, but the exponent may vary from stage to stage. Additionally, the rate of energy expenditure for maintenance, *B*(*m*(*t*))*m*(*t*) in our model, may also not have a simple time-independent prefactor *B*(*m*(*t*)) = *B*_*m*_, since the cost of maintaining a unit of mass can also vary throughout development: in juvenile endothermic birds, before internal heat production has reached its mature level and while there is no energy allocation toward reproductive functions, the maintenance costs are lower than in adult birds [32]. There are also factors that influence metabolic scaling, for example the dimensionality of the environment in which the consumer organism encounters the resources on which it subsists [55]. Another example is a recent model where the partitioning of energy between metabolic processes and heat loss leads to a Π(*m*) that is a linear combination of power law terms with different exponents (2/3 and 1) [29]. Thus it is important to have a general theory, one where Π(*m*(*t*)), *B*(*m*(*t*)) and even *E*(*m*(*t*)) could be arbitrary functions reflecting changes in the organism throughout development. We note that the mathematical formalism developed in the main text, including Eqs. (1-5), is indeed fully general, independent of any specific choice for these functions.

**FIG. S2.**
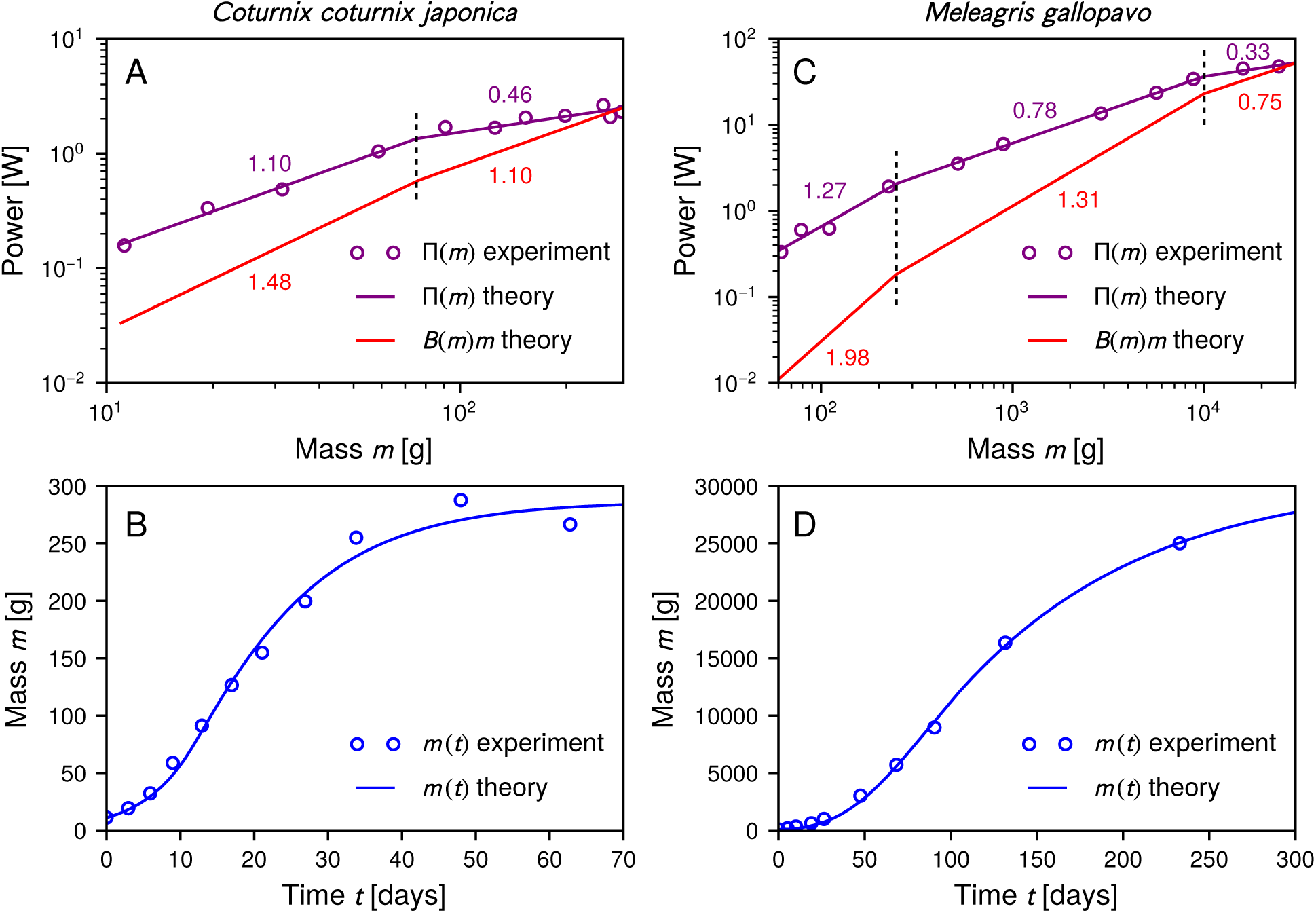
Fits of the generalized growth model to experimental results (taken from Ref. [34]) for male members of two galliform species: Japanese quail (*Coturnix coturnix japonica*, left column); turkey (*Meleagris gallopavo*, right column). A, C: Power input Π(*m*) as a function of mass *m* (purple), with symbols indicating experimental data and lines a piecewise continuous power-law fit to the data. The power *B*(*m*)*m* expended in maintenance is shown in red, and is calculated by fitting the growth model to the mass vs. time data in panel B. Numbers next to the lines indicate the power-law exponents in each regime, while vertical dashed lines show the regime boundaries. B, D: Mean mass *m* versus time *t*, with symbols denoting experimental data and the curve the theoretical best-fit to the generalized growth model.

To illustrate how this formalism can be applied for more complex growth models with varying developmental regimes, we analyze empirical data [34] from two endothermic, galliform bird species: Japanese quail (*Coturnix coturnix japonica*) and turkey (*Meleagris gallopavo*). Fig. S2 A,C shows the experimental resting metabolic rates Π(*m*) (symbols) as a function of *m*. Before applying our selection coefficient theory, we first establish the details of the growth model for each organism. Following the analysis of Ref. [34], we fit Π(*m*) using distinct power-law scaling exponents across different mass regimes (two regimes for quail, three for turkey). The theory fits thus appear as piecewise continuous linear functions (purple lines) on the log-log graphs of panels A and C, with the exponents indicated above each regime. The specific forms of the fitting functions are:

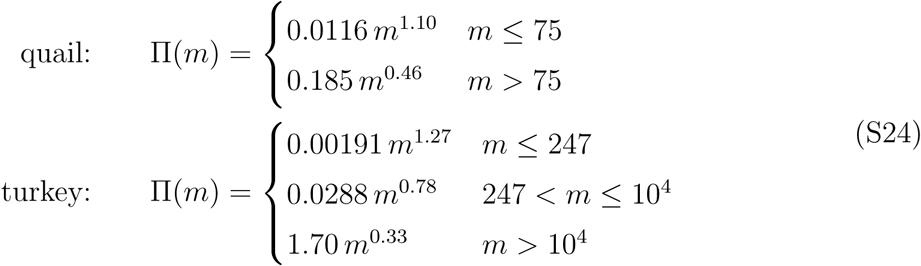

where the units of *m* are g and of Π(*m*) are W. These forms for Π(*m*) fit the experimental data closely, and exhibit a trend seen commonly in organisms with distinct metabolic scaling at different life stages: the power-law exponent progressively decreases as the organism matures (see for example Type III and Type IV scaling behavior as classified by Glazier in Ref. [31]). And though the exponents vary between regimes, they all still fall in the typical biological range ≲ 2.

The resting metabolic power input determines the mass trajectory *m*(*t*) of the organism through the energy conservation equation [Eq. (1) in the main text]:

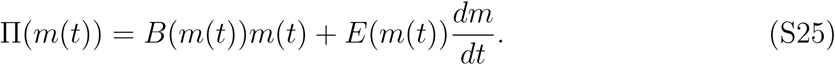

Since we know the empirical *m*(*t*) curves (symbols in Figs. S2 B,D), we can use these to find best-fit forms for *B*(*m*(*t*)) and *E*(*m*(*t*)). For the latter we will assume the cost of synthesizing a new unit mass are constant throughout development, *E*(*m*(*t*)) = *E*_*m*_. This assumption is based on the fact that estimated *E*_*m*_ values are broadly consistent among many different organisms at different life stages, as described in Sec. III above, generally of the order of magnitude *E*_*m*_ ∼ *𝒪*(10^3^) W/g. For *B*(*m*) we allow a more general fitting form, with the product *B*(*m*)*m* (the maintenance power consumption) having piecewise continuous power-law scaling, with the same mass regimes as Π(*m*). Note that in our model *B*(*m*)*m* subsumes all the parts of the resting metabolic power that are consumed in processes other than growth. This is a broader definition of “maintenance” than in models which distinguish the uses of that power, for example separately keeping track of energy used for heat production and reproduction [32]. However for our purposes it is sufficient to collectively track the total non-growth expenditures through *B*(*m*)*m*. The time *t*(*m*) to reach a mass *m*, starting from initial mass *m*_0_ = 11 g (quail), 70 g (turkey), is analogous to Eq. (2) of the main text:

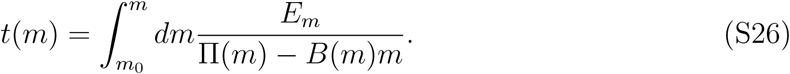

By inverting this to get *m*(*t*) and comparing to the empirical data for the mass trajectories, we can find theoretical best-fits for *E*_*m*_ and the functional forms of *B*(*m*)*m*. The results are *E*_*m*_ = 6005 J/g (quail), 6413 J/g (turkey) and

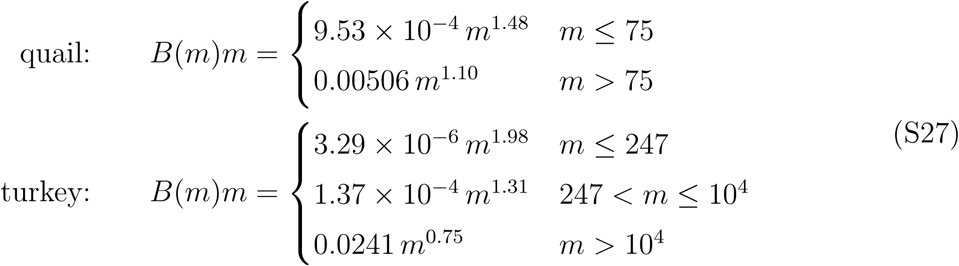

where *B*(*m*)*m* has units of W and *m* has units of g. The fitted theory results (solid curves in Fig. S2 B,D) agree very well with the *m*(*t*) experimental data. The fitted values of *E*_*m*_ are also consistent with the range (1400 − 7500 J/g) seen for various juvenile bird species in an earlier analysis [24]. As expected, the fraction of power input available for growth, 1 − *B*(*m*)*m/*Π(*m*), decreases as the organism develops, eventually reaching zero at the asymptotic adult mass where *B*(*m*)*m* intersects Π(*m*).

With the details of the growth model established, evaluating our expressions for *σ*_*E*_ and *σ*_*B*_ [Eq. (3) of the main text] and comparing them to *σ ′*_*E*_ and *σ′*_*B*_ [Eq. (4) of the main text] is relatively straightforward. Since *E*(*m*(*t*)) = *E*_*m*_ is time-independent, *σ*_*E*_ = *σ′*_*E*_ = 1. This leaves only *σ*_*B*_ and *σ′*_*B*_, given by:

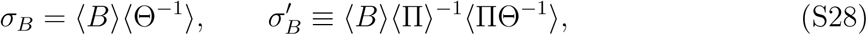

where the averages are given by the following integrals:

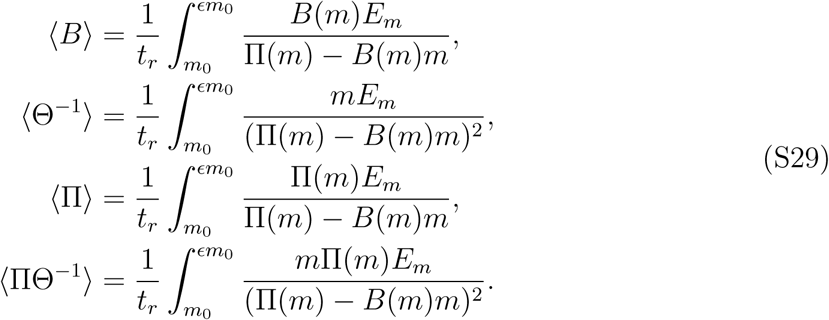

The time for reproductive maturity is *t*_*r*_ = 52 days (quail), 365 days (turkey), based on mean values from the AnAge online database [68]. From the theoretically fitted *m*(*t*) curves this translates to *m*(*t*_*r*_) = *ϵm*_0_ where *E* = 25 (quail), 416 (turkey). The integrals in Eq. (S29) can be evaluated numerically, since all the expressions in the integrand are known based on the growth model fitting. The results for the prefactors in Eq. (S28) are:

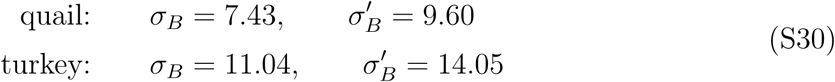

The discrepancy |1 − *σ′*_*B*_*/σ*_*B*_| = 0.27 − 0.29 in both cases. This is comparable to the simple allometric growth model (single power-law) cases investigated in the main text, where the discrepancy was always less than 50%. Thus the relation *s*_*c*_ ≈ − ln(*R*_*b*_)*δC*_*T*_ */C*_*T*_ continues to hold even for more complex growth models in organisms where metabolic scaling varies with developmental stage.

### VI. Generalizing the *s*_*c*_ and 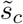 relation for density-dependent population models

The derivation in Sec. I above, relating the baseline selection coefficient *s*_*c*_ to the fractional change in growth rate, 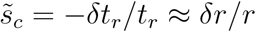, can be generalized to cases where the population growth of the wild-type and mutant organisms is density-dependent. This can occur for example when there is competition for a limited resource shared between the wild-type and mutant (i.e. a nutrient in the case of bacterial growth, or a prey population for a predatory organism), or when there are other external constraints on growth as the overall population increases. As a result, the reproductive time *t*_*r*_ may change from generation to generation, for example lengthening as the resource is depleted and organismal growth is slowed. We will denote 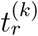 to be the mean reproductive time (duration) of the *k*th generation, and the cumulative time span of *n* generations as 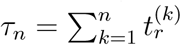, with *τ*_0_ ≡ 0. Eq. (S5), defining the per-generation selection coefficient, can be adapted to this scenario as:

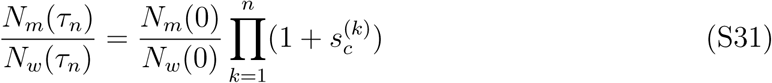

where we have introduced a baseline selection coefficient 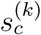 for the *k*th generation that can in general vary from each generation to the next. Eq. (S31) implies

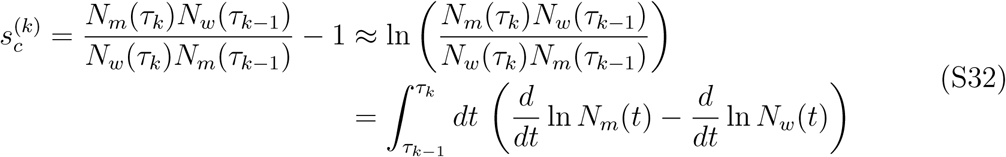

where the approximation assumes selection coefficients 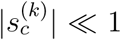, the typical case we consider. The integral expression for 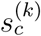 will be useful for evaluating the baseline selection coefficient later. To make further progress, we will define a general density-dependent growth model, and subsequently illustrate it with several examples.

Let us consider wild type and mutant populations of organisms that depend on a shared resource whose quantity *S*(*t*) varies in time. Two examples where this occurs, discussed below, are bacterial populations competing in a chemostat [65–67], and predators competing for the same prey species [60, 62, 64]. The resting metabolic power input has a general functional form Π(*m*(*t*); *S*(*t*), *N*_tot_(*t*)), which depends on the current mass *m*(*t*) of the organism, the amount of available resource *S*(*t*), and potentially also on the total population *N*_tot_(*t*) = *N*_*w*_(*t*) + *N*_*m*_(*t*), i.e if there is a cooperative feeding interaction or other ecological mechanism leading to an Allee effect [59]. Our focus will be on time scales covering many generations, and we will assume changes in Π(*m*(*t*); *S*(*t*), *N*_tot_(*t*)) over a single generation are small enough that we can approximate Π(*m*(*t*); *S*(*t*), *N*_tot_(*t*)) ≈ Π(*m*(*t*); *S*(*τ*_*k*−1_), *N*_tot_(*τ*_*k*−1_)) for *τ*_*k*−1_ *≤ t ≤ τ*_*k*_. Then from Eq. (2) of the main text we can write the reproductive time of the *k*th generation as

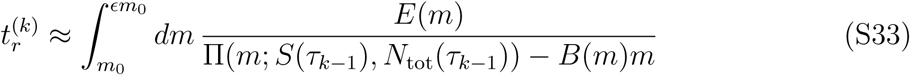

for a wild-type organism with synthesis costs *E*(*m*) and per-mass maintenance costs *B*(*m*). In analogy to the discussion in Sec. I, the corresponding birth rate for the *k*th generation is 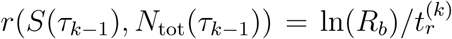, which now depends on *S*(*τk*−1) and *N*_tot_(*τk*−1). To model the population dynamics over time scales much longer than a generation, we make the usual continuum time approximation, *r*(*S*(*τ*_*k*−1_), *N*_tot_(*τ*_*k*−1_)) → *r*(*S*(*t*), *N*_tot_(*t*)), and posit a population model for *N*_*w*_(*t*) of the form

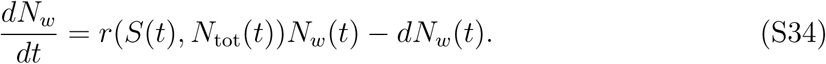

Here *d* is the rate at which the wild-type population is removed from the system i.e. dilution by outflow of solution in a chemostat, or death. If *r* was time-independent, the dynamics of *N*_*w*_(*t*) described by Eq. (S34) would reduce to Eq. (S2) in Sec. I.

We can derive an analogous equation for the mutant population *N*_*m*_(*t*). Under our baseline assumption about the mutant organism, it has modified synthesis costs 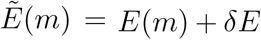 and maintenance costs 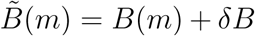, and hence through the analogue of Eq. (S33) it will have mean generation times 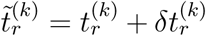 perturbed by 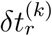 relative to the wild type. In the continuum time population model, this translates to a mutant birth rate 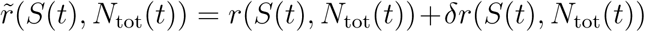, modified by some term *δr*(*S*(*t*), *N*_tot_(*t*)), and population dynamics governed by

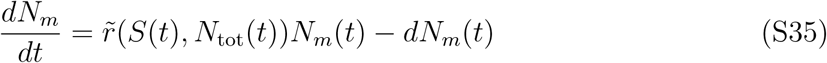

Here *d* is the same as in Eq. (S34), since we assume the mutant is removed at the same rate through dilution or other processes as the wild type. As in the wild type case, for time-independent *r* the solution of Eq. (S35) for *N*_*m*_(*t*) yields the corresponding Sec. I result, Eq. (S3).

The final equation necessary to complete the description of the system is for the resource *S*(*t*). To derive it, first consider the relationship between resource consumption and metabolic expenditure. Over the course of generation *k*, the mean resting metabolic expenditure of a wild type cell per unit time (the average input power) is given by

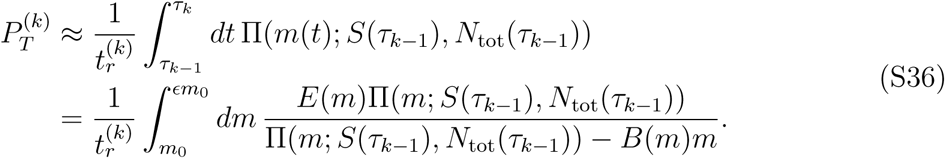

Since this is the generalization of ⟨Π ⟩ from the main text, the corresponding total resting metabolic expenditure 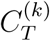 in the *k*th generation is 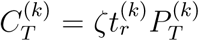. The mean amount of resource consumed per unit time scales with the mean input power as 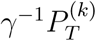, where the yield parameter *γ* sets the conversion rate (flux of resource needed to sustain a certain power input). Since 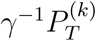 depends on the substrate quantity and total population, let us denote it as a function 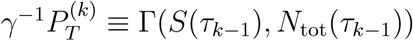, the rate of resource consumption per wild type cell. If the analogous rate for the mutant is 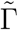, then the equation for *S*(*t*) takes the following form in the continuum approximation,

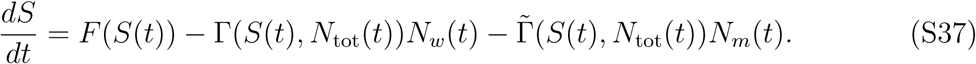

The function *F* (*S*) describes the net production rate of the resource.

From Eqs. (S32), (S34)-(S35) we find:

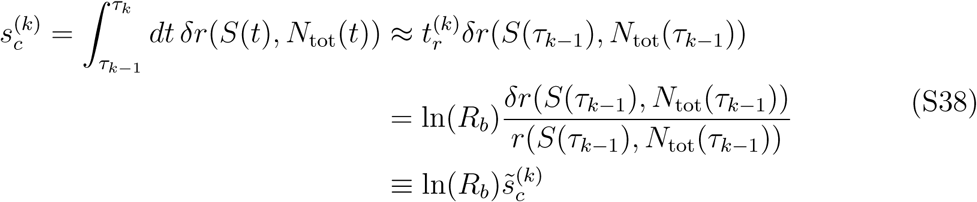

where we have used the fact that 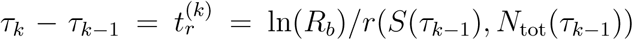, and approximated the integral by assuming *δr*(*S*(*t*), *N*_tot_(*t*)) varies slowly within a generation. We thus have a result directly analogous to the 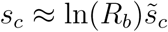 relation derived in Sec. I, but now due to density dependence the fractional change in growth rate 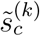 can in general be different for each generation *k*.

It is interesting to note that there are scenarios where 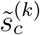 becomes independent of *k*. One case is when the synthesis cost is assumed independent of the organism’s mass, *E*(*m*) = *E*_*m*_, and the maintenance term is negligible *B*(*m*) ≈ 0. This would be a rough approximation for fast-dividing organisms like bacteria. Using the fact that 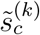 can also be written as 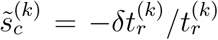, we can see from Eq. (S33) that a mutant with synthesis cost *E*_*m*_ + *δE* would correspond to 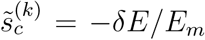. In the notation of the main text, this would mean *σ*_*E*_ = 1, and *σ*_*B*_ is not defined in this case because maintenance is ignored. Note that the *σ*_*E*_ result here is independent of generation number *k*, or any specific details of the density dependence.

Another scenario where 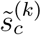 becomes independent of *k* is in the long time limit for cases where the dynamical system of Eqs. (S34)-(S35),(S37) converges to a stationary solution with non-zero total population. In other words *N*_tot_(*t*) and *S*(*t*) go to nonzero asymptotic values as *t* → ∞, which we denote at 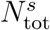 and *S*^*s*^. This requires that *r*_max_, the maximum possible value of *r*(*S, N*_tot_), satisfies *r*_max_ > *d*, and in addition *F* (*S*^*s*^) > 0. Combinining Eqs. (S34)-(S35) we know that

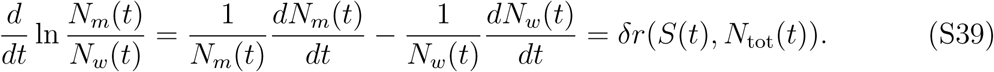

After an initial transient, the right-hand side relaxes to a constant value 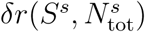. For small *δr*, the regime of interest in our work, we see that the ratio *N*_*m*_(*t*)*/N*_*w*_(*t*) changes very slowly at long times. The system has two possible outcomes: for *δr* > 0 the ratio *N*_*m*_(*t*)*/N*_*w*_(*t*) goes to infinity, corresponding to the mutant eventually taking over the whole population, 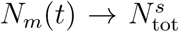. The stationary state values 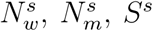, satisfy relations that make the left-hand sides of Eqs. (S34)-(S35),(S37) zero:

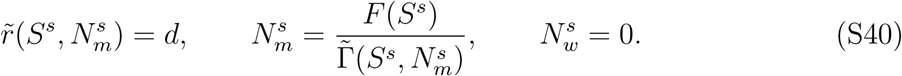

Alternatively when *δr <* 0, *N*_*m*_(*t*)*/N*_*w*_(*t*) goes to zero, with the wild type taking over,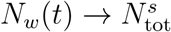. Here the stationary state values satisfy:

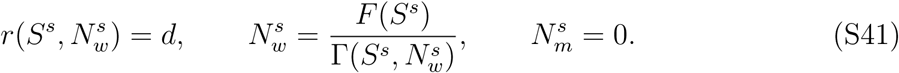

From Eq. (S38) the asymptotic relaxation dynamics at long times between mutant and wild type populations have a selection coefficient proportional to

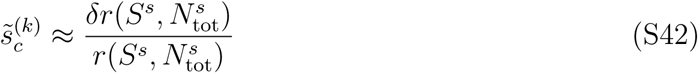

for large *k*. Note that Eq. (S42) also applies if we start in the stationary state with a fully wild type population, and subsequently a mutant with small *δr* appears in the population.

To illustrate the utility of Eq. (S42), it is instructive to consider two model systems:

1) *Bacteria in a chemostat*: The canonical model of a bacterial populations growing in a chemostat [67] assumes resting metabolic power input Π(*m*; *S*; *N*_tot_) takes the form Π(*m*; *S*; *N*_tot_) = Π_max_(*m*)*S/*(*K*_*s*_ + *S*), which depends on a resource *S* but not explicitly on *N*_tot_. Here Π_max_(*m*) is the power input under unlimited resource, and the second term is a phenomenological hyperbolic function, first posited by Monod [63], that expresses approximately linear dependence of the power on resource for small *S*, and saturation for *S* much greater than some threshold value *K*_*s*_. We assume a simple allometric growth model with exponent *α* = 1 typical of bacteria: Π_max_(*m*) = Π_0_*m* for some constant Π_0_, *E*(*m*) = *E*_*m*_, *B*(*m*) = *B*_*m*_. The integrals in Eq. (S33) for 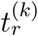 and Eq. (S36) for 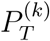 can be evaluated directly, and the resulting birth rate *r*(*S*(*t*)) and resource consumption rate Γ(*S*(*t*)) take the form

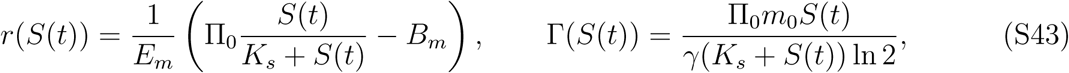

where we have taken *R*_*b*_ = *ϵ* = 2 for simplicity. (Other values of *ϵ* would just add an overall prefactor.) If *B*_*m*_ is neglected, a common simplification for bacteria, *r*(*S*) takes the form known as Monod’s law, *r*(*S*) = *r*_max_*S/*(*K*_*s*_ + *S*) with a maximum rate *r*_max_ = Π_0_*/E*_*m*_. However for completeness we will not ignore *B*_*m*_, and use Eq. (S43) instead. For typical chemostat experiments, cell death is negligible relative to *D*, the dilution rate. This is the rate at which the solution (including the resource and bacterial cells) is removed from the system. Hence we set *d* = *D*. The final piece of the chemostat model is the net resource production, *F* (*S*) = *F*_0_ − *DS*, where *F*_0_ represents the incoming resource from a reservoir, and −*DS* is the output flow of resource due to dilution. The dilution rate in chemostat experiments is typically of the order of *D* ∼ *𝒪*(0.1) hr^−1^ [79].

Given these model details, Eqs. (S34)-(S35),(S37) have a a stable stationary solution with 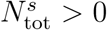 in the long time limit if *D <* (Π_0_ − *B*_*m*_)*/E*_*m*_. For concreteness let us take *δr <* 0, so the stationary state satisfies Eq. (S41). Using Eq. (S43) to calculate *δr* for perturbations to *E*_*m*_ and *B*_*m*_, we plug into Eq. (S42) in the stationary limit to get

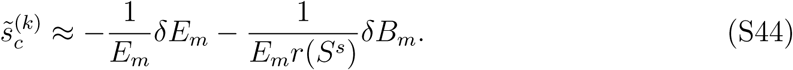

Hence

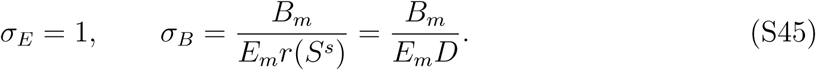

We thus see that the relative maintenance contribution *σ*_*B*_ to the baseline selection coefficient can be tuned by changing *D*, making the chemostat a versatile experimental tool to explore maintenance versus synthesis cost selection pressures in bacterial populations. As discussed in Sec. III above, empirical data for a broad swath of unicellular organisms [12] yields a global estimate *B*_*m*_*/E*_*m*_ = 3 *×* 10^−6^ s^−1^. For a typical experimental range *D* = 0.05 − 0.5 hr^−1^, we then know that *σ*_*B*_ would vary between 0.22 and 0.022.

As noted above, the total resting metabolic expenditure in the *k*th generation 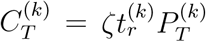, which can be written as 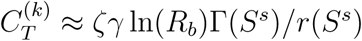 for large *k* when the system approaches the stationary state. Using Eq. (S43) to calculate 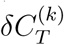 for perturbations to *E*_*m*_ and *B*_*m*_, we find

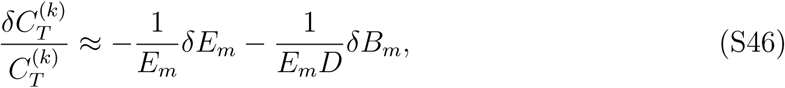

and so

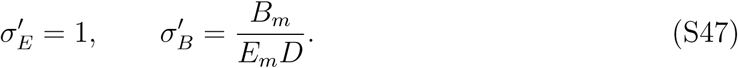

Comparing Eqs. (S45) and (S47), we see that *σ*_*E*_ = *σ′*_*E*_, *σ*_*B*_ = *σ′*_*B*_. Thus the approximation 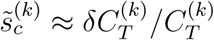 works well in this stationary limit.

2) *Predator-prey dynamics*: The second example is predator-prey system where the predator wild type organism of population *N*_*w*_ has a resting metabolic rate described by some allometric exponent *α*, so that Π(*m*; *S*) = Π_0_(*S*)(*m/m*_0_)^*α*^, with a function Π_0_(*S*) that depends on the population of a prey organism *S*. For convenience we have explicitly factored out 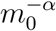, in order that Π_0_(*S*) have units of W. As in the allometric model of the main text, *E*(*m*) = *E*_*m*_, *B*(*m*) = *B*_*m*_. From Eq. (S33) the predator reproductive time for *α* ≠ 1 is given by:

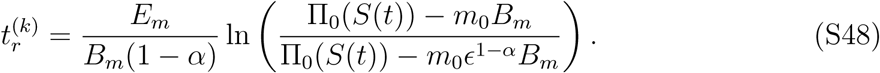

The population growth rate 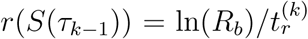. Since the exponent *α* for metazoans is typically close to 1 (as in the common choice of *α* = 3*/*4 [23, 24]), we will simplify the expression for *r*(*S*(*τ*_*k*−1_)) by expanding it to first order in *α* around *α* = 1. In the continuum version, where *S*(*τ*_*k*−1_) → *S*(*t*), the growth rate then takes the form

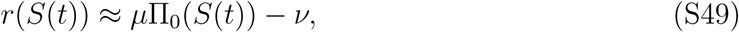

where

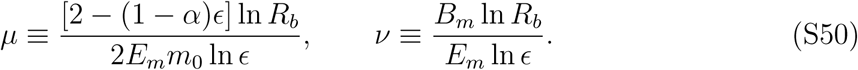

This approximation works best when |1 − *α*| ln *ϵ* ≪ 2. The first term in the *r*(*S*(*t*)) expression is the contribution to population growth of metabolic power input through prey consumption, while the second term describes the degree to which the growth rate is reduced by having some of that power channeled into maintenance. We approximate Eq. (S36) for 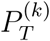 to lowest order in 1 − *α*, giving the following expression for the resource consumption rate,

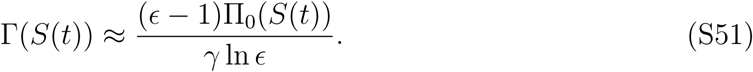

Eqs. (S49)-(S51) can be plugged into the dynamical system of Eqs. (S34)-(S35),(S37). To complete the description of the predator-prey system, we make several additional assumptions: we interpret *d* as the death rate of the predator, and choose a prey reproduction function *F* (*S*) = *bS*(1−*S/K*), where *b* is the maximum net rate at which the prey species reproduces itself, *K* is the prey carrying capacity. Finally we choose 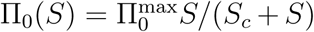 for some constants 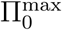 and *S*_*c*_, where 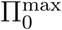 is the maximum metabolic power input in the limit of unlimited prey abundance, and *S*_*c*_ is the critical prey population above which saturation of Π_0_(*S*) occurs. This has the form of a Holling type II predator-prey functional response [61, 62], which assumes that when prey becomes overabundant, the consumption rate is limited by the finite time required for the predator to process each kill, hence leading to saturation of Π_0_(*S*). Given the above assumptions, the dynamical system of Eqs. (S34), (S37) reduces to the Rosenzweig-Macarthur (RM) predator-prey model [60, 64] in the absence of a mutant predator population, and with maintenance *B*_*m*_ neglected. Here we consider the generalized RM model with *B*_*m*_ > 0 and with populations of both wild type and mutant predators competing for the same prey.

To apply Eq. (S42), we need to establish the properties of the stationary solution of Eqs. (S34)-(S35),(S37), along the lines of the stability analysis for the original RM model in Ref. [60]. For concreteness let us assume *δr <* 0, so the stationary solution should satisfy Eq. (S41). Then a stationary solution exists with 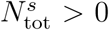 if 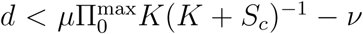, and takes the form

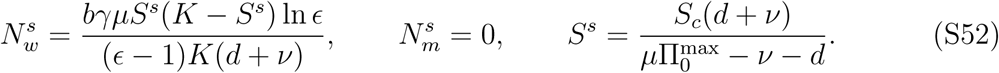

This solution is stable when *S*^*s*^ > (*K* − *S*_*c*_)*/*2, which will be the parameter regime on which we focus. When the stationary solution is unstable, the system exhibits limit cycles, which are beyond the scope of this analysis (though in such cases Eq. (S38) would still hold, with an 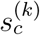 that becomes periodic in *k* in the long time limit).

Putting everything together, we can use Eqs. (S49)-(S50) to calculate *δr* for perturbations to *E*_*m*_ and *B*_*m*_, and we then evaluate Eq. (S42) at the stationary state,

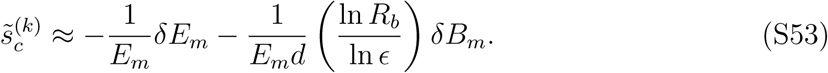

Thus in this case,

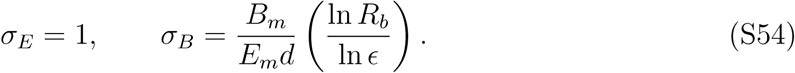

The final result for *σ*_*B*_ has a very similar form to the bacterial chemostat example above, Eq. (S45), except for an additional factor of ln *R*_*b*_*/* ln *E*, and *d* being the predator death rate, rather than a dilution rate. As discussed in Sec. III above, the ratio *B*_*m*_*/E*_*m*_ is expected to be somewhat different for multicellular versus unicellular species, with for example *B*_*m*_*/E*_*m*_ = 10^−7^ − 10^−6^ s^−1^ for a range of mammals [24]. The scale of the dimensionless factor ln *R*_*b*_*/* ln *E* can also be estimated. For example, data from a variety of fissiped carnivore and insectivore mammals yields a range ln *R*_*b*_*/* ln *ϵ* ≈ 0.2 − 0.4 [58]. Given these estimates, we will get *σ*_*B*_ > *σ*_*E*_ = 1 when the mean predator lifetime *d*^−1^ is on the order of a year or higher. If we do the analogous 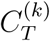 perturbation analysis as in the previous example, we again find that *σ*_*E*_ = *σ*^*′*^_*E*_, *σ*_*B*_ = *σ*^*′*^_*B*_, validating the approximation 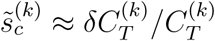 for large *k*.

### VII. Understanding the roles of baseline metabolic costs versus adaptive effects in selection

In the main text our focus has been on understanding the baseline contribution to selection, *s*_*c*_, arising from changes in metabolic costs (perturbations to synthesis and maintenance). However in the general case a genetic variant will have additional perturbations to its phenotype contributing to the selection coefficient, yielding an overall coefficient *s* = *s*_*c*_ + *s*_*a*_ with an adaptive correction *s*_*a*_ [12]. In this section we illustrate how our bioenergetic growth formalism can be extended to consider perturbations beyond metabolic costs, yielding expressions for *s*_*a*_. We show that even with these additional perturbations, *s*_*c*_ retains the form of our original formalism, and can be generally related to associated fractional changes in the total resting metabolic expenditure *C*_*T*_. However the relationship between *s*_*a*_ and *C*_*T*_ is more complicated, dependent on system-specific details of the organism and its environment. Thus investigating the relative magnitude of *s*_*c*_ and *s*_*a*_ in individual biological cases, and the ways in which *s*_*a*_ might or might not compensate for the metabolic costs encapsulated in *s*_*c*_, opens a rich set of questions for future study.

We proceed by generalizing the derivations in SI Secs. and to include two examples of adaptive effects for the mutant organisms: changes in the death rate *d*, and changes in resting metabolic power input Π(*m*(*t*)). These are not the only quantities in the theory that could potentially be subject to adaptation: other possibilities include the the mean number of offspring *R*_*b*_ or the reproductive maturity parameter *ϵ*. However these two examples illustrate the general scheme for how to include adaptive effects in the formalism. A more comprehensive treatment of the interplay between baseline metabolic costs and adaptive effects in selection is the subject of an upcoming work [51].

In the original derivation of SI Sec. I, where we considered only the baseline selective contribution due to metabolic costs, we assumed the mutant has the same mean number of offspring *R*_*b*_ and death rate *d* = − ln(*p*(*t*_*r*_))*/t*_*r*_ as the wild type. We now partially relax that assumption: we still keep *R*_*b*_ the same, but allow the mutant death rate 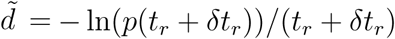 to be different than *d* by some amount 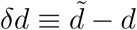. For example if the survival probability *p*(*t*) was exponential for the wild-type, *p*(*t*) = exp(−*ηt*), and the rate *η* was modified in the mutant to 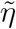, then 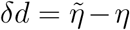. One case where such considerations would come into play would be bacterial strains growing in an environment with antibiotics: a variant that acquired antibiotic resistance would have a lowered death rate, *δd <* 0, but also possibly non-trivial metabolic costs associated with maintaining the resistance [52– 54], as discussed in more detail below. Since the mutant now has both altered birth rate 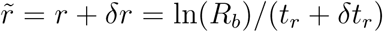 and death rate 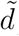, the population growth equation [Eq. (S3)] becomes

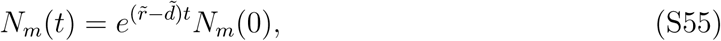

and the ratio of mutant to wild-type populations [Eq. (S4)] is changed to

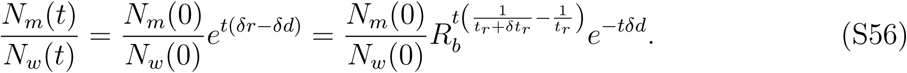

Since we now have both metabolic costs and adaptive effects, Eq. (S5) now defines the full selection coefficient *s* instead of just *s*_*c*_, evaluated after *n* wild-type generations, *t* = *nt*_*r*_,

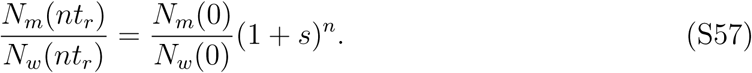

Comparing Eq. (S57) to Eq. (S56), we can write

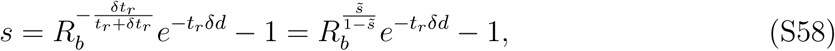

where 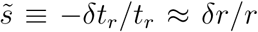, with the latter approximation valid when 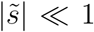. Note that in this general case *δr* (or equivalently *δt*_*r*_) describes the perturbation to the growth rate both from the altered metabolic costs as well as adaptive effects (as we will see below when we consider changes in Π(*m*(*t*))). We will distinguish the metabolic versus adaptive contributions shortly. If the magnitudes of *δd* and *δt*_*r*_ are small relative to *t*_*r*_, Eq. (S58) to first order in the perturbations *δd* and *δt*_*r*_ is given by

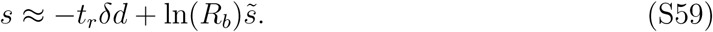

To complete the description of *s*, we now would like to expand out the 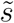 contribution on the right-hand side of Eq. (S59) into contributions from metabolic and adaptive perturbations, generalizing the derivation of SI Sec. II for 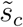. Starting with Eq. (S9) for *t*_*r*_, we consider how *t*_*r*_ will be altered under the simultaneous perturbations: *E*(*m*) → *E*(*m*) + *δE, B*(*m*) → *B*(*m*) + *δB*, Π(*m*) → Π(*m*) + *δ*Π(*m*). The first two are the baseline metabolic changes in synthesis and maintenance costs we considered before. The last one represents the possibility that the genetic variant could also have modifications in the resting metabolic power input function Π(*m*(*t*)). As before we assume *δE* and *δB* are independent of *m*(*t*) for simplicity, but we allow the power input perturbation *δ*Π(*m*(*t*)) to have some general *m*(*t*)-dependent functional form. Eq. (S10) then becomes

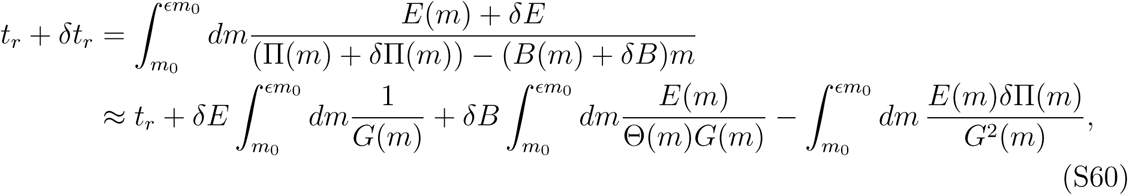

where we have expanded to first order in the perturbations, and *G*(*m*) = Π(*m*) − *B*(*m*)*m*, Θ(*m*) = *G*(*m*)*/m*. As in SI Sec. II, we can rewrite the perturbation terms on the right-hand side of Eq. (S60) in terms of averages over 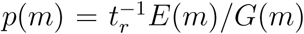, and get an expression for 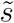,

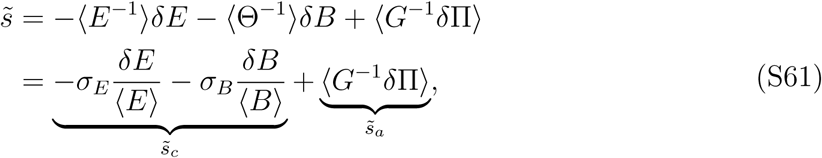

where the synthesis/maintenance prefactors have the same definition as before: *σ*_*E*_ = ⟨*E*⟩⟨*E*^−1^⟩, *σ*_*B*_ = ⟨*B*⟩⟨Θ^−1^⟩. The first two terms of Eq. (S61) are thus 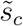 [Eq. (S13)], the baseline contribution to 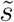 from perturbations to metabolic synthesis/maintenance costs. The new adaptive contribution 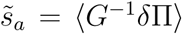 comes from the perturbation to the power input, and has a straightforward physical interpretation: since *G*(*m*) is the power available for growth (once maintenance costs are subtracted away from the power input), the average ⟨*G*^−1^*δ*Π⟩ is just the fractional size of the power input perturbation *δ*Π(*m*) relative to *G*(*m*), averaged over the course of a generation. Plugging Eq. (S61) into Eq. (S59), we get our final expression for *s*,

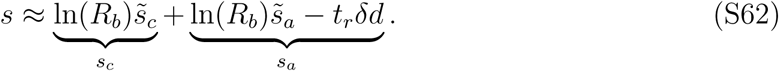

The first term is *s*_*c*_, what we had calculated originally under the baseline assumption, and the remaining terms we can identify as *s*_*a*_, the correction due to adaptive effects beyond the baseline assumption.

As we did for the baseline case, we can compare the contributions to 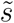 in Eq. (S61) from the different perturbations to the associated changes in the total resting metabolic expenditure per generation, *C*_*T*_ = *ζt*_*r*_(Π). Again proceeding analogously to SI Sec. II, Eqs. (S15)-(S19), we find the fractional change

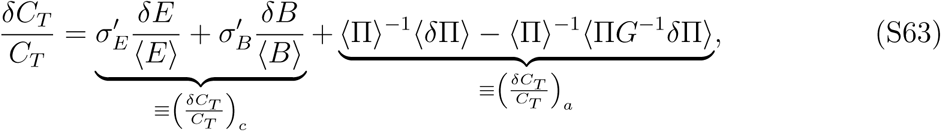

where we have split *δC*_*T*_ */C*_*T*_ into the baseline cost contribution (*δC*_*T*_ */C*_*T*_)_*c*_ and the adaptive correction (*δC*_*T*_ */C*_*T*_)_*a*_. Here *σ′*_*E*_, *σ′*_*B*_ have the same definitions as before, given in Eq. (S19). As we argued in the main text, the prefactors *σ′*_*E*_, *σ′*_*B*_ are comparable to *σ*_*E*_, *σ*_*B*_ respectively across a wide range of biologically relevant growth scenarios. Thus the maintenance/synthesis contributions to 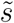 in Eq. (S61) have a direct reflection in the baseline contribution to *δC*_*T*_ */C*_*T*_ in Eq. (S63). In other words 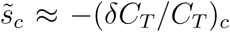, the baseline relation that is the central focus of the main text.

For the adaptive components, the relation between 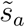 from Eq. (S61) and (*δC*_*T*_ */C*_*T*_)_*a*_ from Eq. (S63) is not so straightforward. Moreover the full adaptive contribution to the selection coefficient, *s*_*a*_ from Eq. (S62), includes a term reflecting the change in death rate, −*t*_*r*_*δd*, which has no analogue in (*δC*_*T*_ */C*_*T*_)_*a*_. As a concrete example, consider the allometric growth model described in the main text, where Π(*m*(*t*)) = Π_0_*m*^*α*^(*t*), *E*(*m*(*t*)) = *E*_*m*_, *B*(*m*(*t*)) = *B*_*m*_. Let us focus on the simplest case, *α* = 1, corresponding to exponential cell mass growth, where the baseline relation is exact, 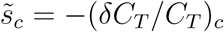 since *σ*_*E*_ = *σ′*_*E*_ = 1 and *σ*_*B*_ = *σ′*_*B*_. If the power input perturbation occurs through the prefactor, Π_0_ → Π_0_ + *δ*Π_0_, then *δ*Π(*m*) = *δ*Π_0_*m*. We can then evaluate both 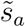 and (*δC*_*T*_ */C*_*T*_)_*a*_:

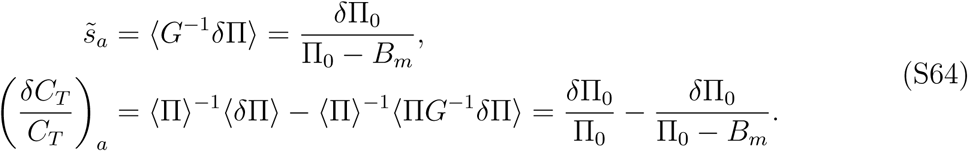

So we see that even in this simple case 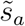 and (*δC*_*T*_ */C*_*T*_)_*a*_ differ by an extra term in the latter. The significance of this difference depends on the details of the specific system under consideration. A schematic summary of the above results is shown in Fig. S3.

The relative contributions of *s*_*c*_ and *s*_*a*_ to selection in particular biological systems may also depend on aspects of the environment, with *s*_*c*_ or *s*_*a*_ playing a leading role in different contexts. For bacterial strains evolving in the presence of an antibiotic, the adaptive benefit of a mutation that confers antibiotic resistance (i.e. lowering the death rate *d* in the example above) will be reflected in a significant positive fitness contribution *s*_*a*_ > 0. But the molecular mechanisms that lead to resistance typically come with non-negligible metabolic costs (*s*_*c*_ *<* 0) that slow bacterial growth in the absence of drug, for example over-expression of energy-consuming multi-drug efflux pumps [50], or switching to less energy-efficient biochemical pathways that are not targeted by the drug [52]. While *s*_*a*_ may more than compensate for *s*_*c*_ when the antibiotic is present, the resistant strains are at least initially at a disadvantage in environments without the antibiotic, where *s*_*c*_ is likely the dominant contribution to *s*. Ref. [54] conducted meta-analysis of competitive fitness assays between wild type and resistant bacteria in drug-free conditions, focusing on cases where the resistance was conferred by a single mutation. They found mean values of *s* for mutants resistant to different drugs to range between ≈ −0.03 to −0.28 for eight different bacterial species. Given the large effective populations of bacteria, these values are large enough to put the mutant strains under strong selective pressure to reduce the metabolic price of maintaining resistance. Intriguingly the net result of this pressure under subsequent drug-free evolution is typically not the loss of the resistance, but rather additional mutations that compensate for the metabolic costs [52–54].

**FIG. S3.**
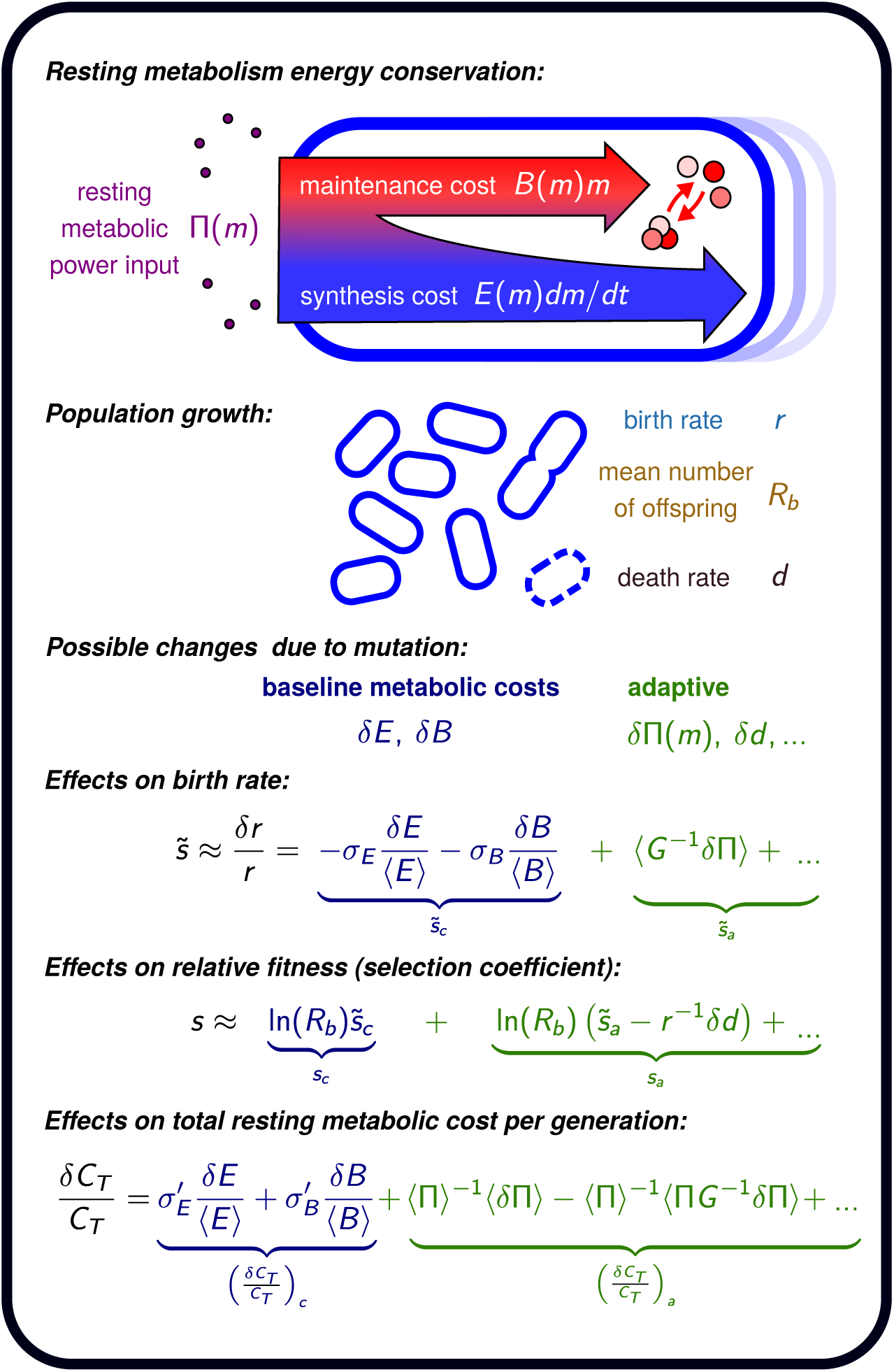
Schematic summary of the basic theoretical formalism. Energy conservation for individual organisms implies a growth relation for mass *m*(*t*): resting metabolic power input Π(*m*) is expended in maintenance *B*(*m*)*m* and synthesis *E*(*m*)*dm/dt* [main text Eq. (1)]. The overall population has a birth rate *r* = ln(*R*_*b*_)*/t*_*r*_ and death rate *d*, where *R*_*b*_ is the mean number of offspring and *t*_*r*_ is the mean generation time [related to metabolism through main text Eq. (2)]. Mutations can introduce perturbations in the baseline metabolic costs *δE, δB* as well as adaptive effects like *δ*Π(*m*), *δd*. These have several consequences: i) modifying the generation time by *δt*_*r*_ [SI Eq. 60], which in turn changes the birth rate by *δr* and leads to a relative fitness difference quantified by the selection coefficient *s*; ii) modifying the total resting metabolic expenditure *C*_*T*_ over a generation by an amount *δC*_*T*_. All these changes can be expressed in terms of time averages of metabolic quantities integrated over the wild type generation time *t*_*r*_, i.e. 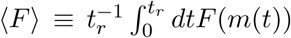. The definitions of *σ*_*E*_, *σ*_*B*_, *σ′*_*E*_, *σ′*_*B*_ are given in main text Eqs. (3-4), and *G*(*m*) ≡ Π(*m*) − *B*(*m*)*m ≥* 0 is the power channeled to growth. The baseline metabolic contributions due to *δE* and *δB* we denote with subscript *c*, and the remaining (adaptive) contributions with subscript *a*.

Another scenario where environmental conditions can determine the significance of metabolic costs to selection is for systems that exhibit spare respiratory capacity: an energetic reserve that allows them to maintain ATP levels to cope with stress that increases energy demand [42, 43, 45, 46]. While this is particularly relevant for multicellular eukaryotes, it may also be facilitated in prokaryotic populations by large cell-to-cell variations in ATP concentrations [44]. If such spare capacity exists, then under unstressed conditions a mutation with extra baseline metabolic costs may get absorbed by the reserve, with these costs not translating into a penalty in terms of cell growth rate (up to a certain cost threshold). In other words, after factoring in the spare capacity the effective *δE* and *δB* of this mutation would have a smaller magnitude (or be zero) compared to the case where no spare capacity existed, and hence the magnitude of *s*_*c*_ would also be smaller. However these costs would not necessarily always remain hidden from selection: if the environment switched to stress conditions with higher energy demands over the long term, these mutants would have a lower reserve capacity to cope. In this case there would be less buffering of the costs, and a larger magnitude of *s*_*c*_ for the mutants.

In summary, quantitative understanding of how *s* splits into *s*_*c*_ and *s*_*a*_ components, and their relative importance under different conditions, will be very useful in the probing the metabolic influences on evolution in biological systems, and an important target for future work [51].

